# HiFAA: a high-confidence framework for transcription factor footprinting with ATAC-seq

**DOI:** 10.1101/2025.10.11.681777

**Authors:** Hijiri Inoue, Chisato Okamoto, Hisakazu Kato, Osamu Tsukamoto, Seiji Takashima, Ken Matsuoka

## Abstract

Accurate inference of transcription factor (TF) binding from chromatin accessibility remains a major challenge, as conventional methods often miss true binding events, yield unstable motif rankings, or fail to resolve dynamic changes. Here we present high-confidence transcription factor footprint analysis using ATAC-seq (HiFAA), a framework that integrates topologically associating domain–restricted analysis, single-base-resolution footprint construction, and footprint ranking based on footprint bottom height. HiFAA markedly improves sensitivity while maintaining high specificity, enabling detection of binding events overlooked by existing analyses. It also ranks truly bound TFs with higher motif scores, enhancing predictive accuracy, and maintains consistently high geometric mean (G-mean) values, providing a stable balance between sensitivity and specificity. Furthermore, its high-resolution footprints capture subtle structural variations, allowing precise detection of developmental stage–specific TF-binding. By uniting sensitivity, specificity, and dynamic resolution, HiFAA establishes a robust paradigm for reliable TF footprinting from ATAC-seq data.

## Introduction

Transcription factors (TFs) are central regulators of gene expression, modulating development, cell fate decisions, and disease progression through selective DNA binding at promoters, enhancers, and other regulatory elements. Precise identification of TF-binding sites is therefore a key objective in regulatory genomics. Typically, chromatin immunoprecipitation followed by sequencing (ChIP-seq) has been the gold standard for mapping TF-binding sites in vivo^1,2^. While powerful, ChIP-seq suffers from critical limitations: reliance on high-quality antibodies that are unavailable for many TFs, large input requirements that preclude rare cell populations, and limited resolution due to chromatin fragmentation. These constraints limit its applicability in contexts where material is scarce, or nucleotide-level precision is needed.

Assay for transposase-accessible chromatin using sequencing (ATAC-seq)^3^ provides a highly sensitive alternative by profiling genome-wide open chromatin regions from small amounts of input material. Because TFs preferentially bind within accessible chromatin, ATAC-seq data have the potential to provide indirect yet powerful insights into TF occupancy. Indeed, TF binding produces footprints—localized depletions of transposase insertion events within accessible regions—that can serve as signatures of direct TF–DNA interactions^4,5^.

Several computational methods have been developed to exploit this principle, yet they differ substantially. HOMER^6^, a widely used motif enrichment tool, identifies TF motifs within accessible peaks but does not consider ATAC-seq signal structure. As such, it achieves high specificity but suffers from low sensitivity. In contrast, TOBIAS^7^ and related footprinting algorithms (e.g., HINT-ATAC^8^) improve sensitivity by identifying valleys of reduced accessibility but rely largely on footprint width or depth, often yielding unstable ranking and false positives. Thus, while HOMER and TOBIAS remain important, each captures only part of the regulatory signal embedded in ATAC-seq data.

To overcome these limitations, we developed HiFAA (High-confidence transcription factor Footprint Analysis using ATAC-seq datasets), a framework designed to deliver both sensitivity and accuracy. HiFAA introduces three key innovations: (i) restricting analysis to the topologically associating domain (TAD) encompassing the target gene^9–11^, which reduces the search space by >90% and enriches for biologically relevant elements; (ii) constructing single-base-resolution footprints using Binary Alignment Map (BAM) data and pseudo-reads, achieving near-perfect concordance with ChIP-seq peak summits5; and (iii) implementing a novel ranking system based on footprint bottom height to distinguish true TF footprints from artifacts.

Together, these innovations yield three advances: HiFAA improves sensitivity while maintaining specificity, ranks truly bound TFs with higher scores, and detects subtle, condition-dependent changes in footprint structure such as stage-specific TF binding. Benchmarking across TFs and tissues confirmed that HiFAA outperforms conventional analyses. Importantly, HiFAA detected dynamic TF-binding events previously inaccessible, including GATA1 occupancy at the BCL11A enhancer during the fetal-to-adult hemoglobin switch^12,13^. By combining structural resolution, biological context, and balanced evaluation, HiFAA substantially extends the capabilities of ATAC-seq–based motif analysis.

## Results

### HiFAA framework and analytical workflow

We developed HiFAA to extract biologically meaningful footprints from ATAC-seq data and distinguish genuine TF-binding events from background signals or false positives. The framework consists of three major steps: (1) restricting analysis to the TAD containing the target gene, (2) constructing high-resolution footprints using BAM data and pseudo-reads, and (3) ranking and selecting footprints based on footprint bottom height (Fig. 1a). This workflow preserves biological context and ensures reliability.

**Figure 1.**
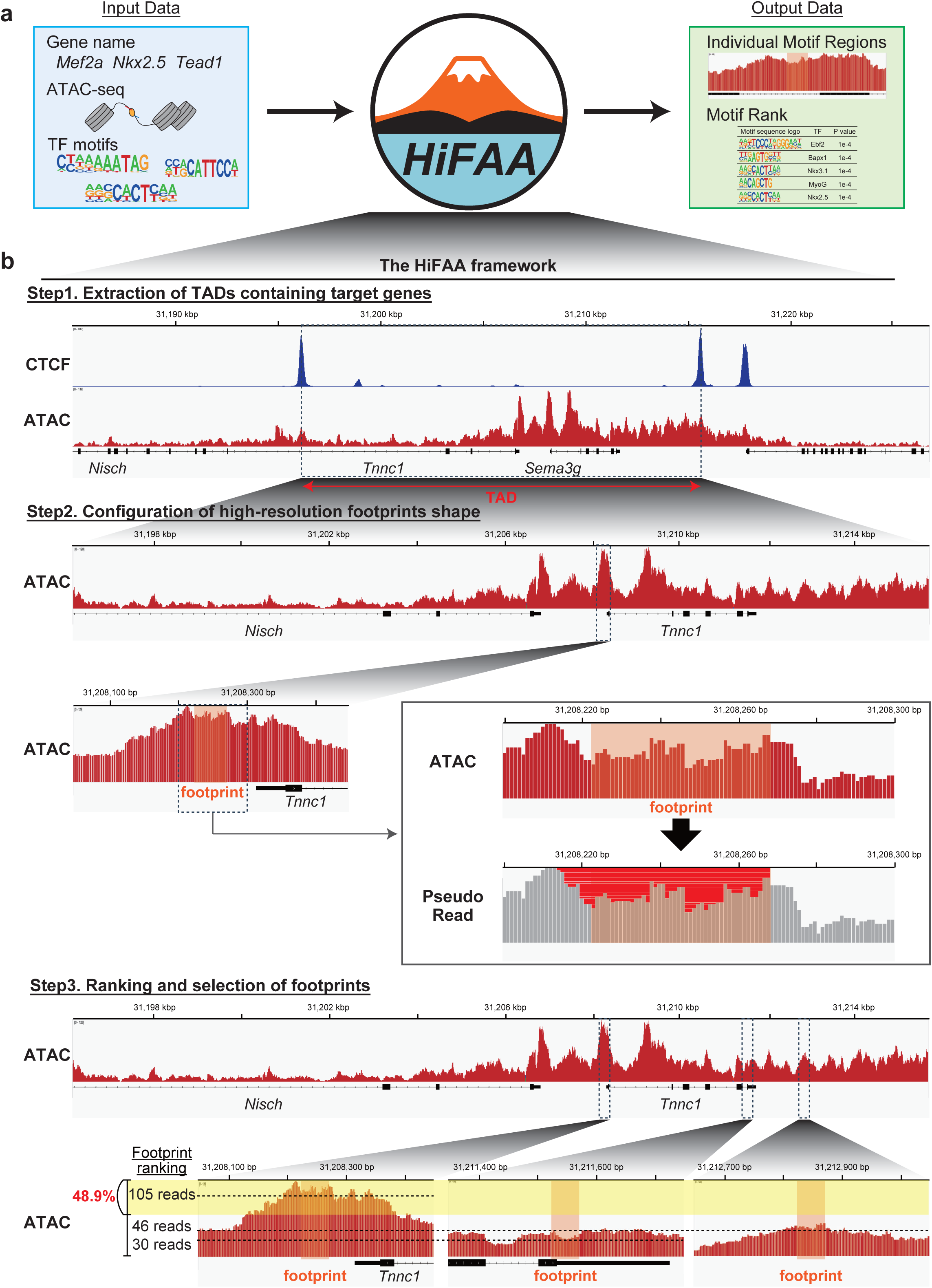
Overview of the HiFAA framework for transcription factor footprint analysis. **a,** Conceptual overview of inputs and outputs. HiFAA integrates user-defined target genes, ATAC-seq data, and known transcription factor (TF) motifs to generate rank-ordered footprints annotated with candidate TF-binding events. **b,** Workflow of HiFAA. Step 1: identification of the topologically associating domain (TAD) containing the target gene and restriction of subsequent analysis to that region. Step 2: reconstruction of high-resolution footprints at single-base resolution using BAM data and pseudo-reads. Step 3: ranking and filtering of footprints based on footprint bottom height, retaining the top 48.9% for optimal specificity and sensitivity.

### Step 1: Restricting analysis to TADs

The first step in the HiFAA workflow is to identify the TAD containing the gene of interest and to restrict motif analysis to this domain (Fig. 1b). TADs represent higher-order chromatin structures that demarcate functional regulatory units in which enhancer–promoter interactions occur with high frequency^1,2,14^. By restricting analysis to the relevant TAD, HiFAA selectively analyzes regulatory regions highly relevant to the target gene, while excluding genomic regions unlikely to contribute to its transcriptional regulation.

This targeted strategy provides several advantages and substantially reduces computational burden by decreasing the size of the sequence space analyzed. Furthermore, restricting the search space reduces the likelihood of identifying spurious footprints in regions unrelated to the gene of interest, thereby lowering false-positive rates. In contrast, genome-wide scanning by methods such as TOBIAS^7^ and HINT-ATAC^8^ increases noise and cost. HiFAA instead emphasizes biologically constrained analysis aligned with three-dimensional genome architecture.

### Step 2: Single-base-resolution footprints from ATAC-seq signal

Traditional motif analysis methods, such as HOMER^6^, infer TF binding by scanning the centers of ATAC-seq peaks identified by MACS2^15^. While this strategy is straightforward, it disregards the fine structure of ATAC-seq read distribution and is prone to false positives when motifs are present in accessible chromatin regions not occupied by TFs. Footprint-based methods, including TOBIAS^7^ and HINT-ATAC^8^, attempt to improve on this by identifying regions with reduced transposase accessibility (footprint “valleys”) flanked by elevated insertion frequencies. Although these approaches improve sensitivity and specificity, they rely primarily on simple metrics such as footprint width or depth and often neglect subtle but biologically important variations in footprint shape.

Our comparative analyses revealed that outer boundaries of footprints are generally stable, but internal structures—particularly bottom shape and height—shift with TF occupancy. Capturing these subtle variations provides sensitivity beyond conventional approaches. We hypothesized that leveraging such internal structural information could provide greater sensitivity for detecting TF-binding events.

In HiFAA Step 2, BAM files are processed to extract high-resolution ATAC-seq signals across peak regions. Pseudo-reads are interpolated between peaks to generate continuous, single-base-resolution profiles of transposase accessibility (Fig. 1b). This approach allows footprints to be defined with unprecedented detail. For each footprint, the signal is measured continuously from the footprint bottom to the summits of adjacent peaks. By capturing subtle variations within footprints, HiFAA detects condition-dependent structural changes overlooked by conventional approaches.

### Improved sensitivity in motif detection

To assess whether the high-resolution footprint construction in HiFAA improves motif detection sensitivity, we performed a direct comparison with HOMER and TOBIAS (Fig. 2a). We focused on three cardiac transcription factors—MEF2A, NKX2.5, and TEAD1—using adult mouse cardiac tissue^16^. For each TF, publicly available ChIP-seq datasets were obtained, and peaks were called using MACS2^15^. The top 100 peaks were extracted, and reproducible peaks common to both replicates were retained, yielding 77 peaks for MEF2A, 36 peaks for NKX2.5, and 70 peaks for TEAD1. In parallel, we generated independent ATAC-seq datasets from adult mouse hearts under matched conditions. These ATAC-seq data were then analyzed with HiFAA, HOMER, and TOBIAS, and motif detection sensitivity was evaluated relative to the reference ChIP-seq peaks.

**Figure 2.**
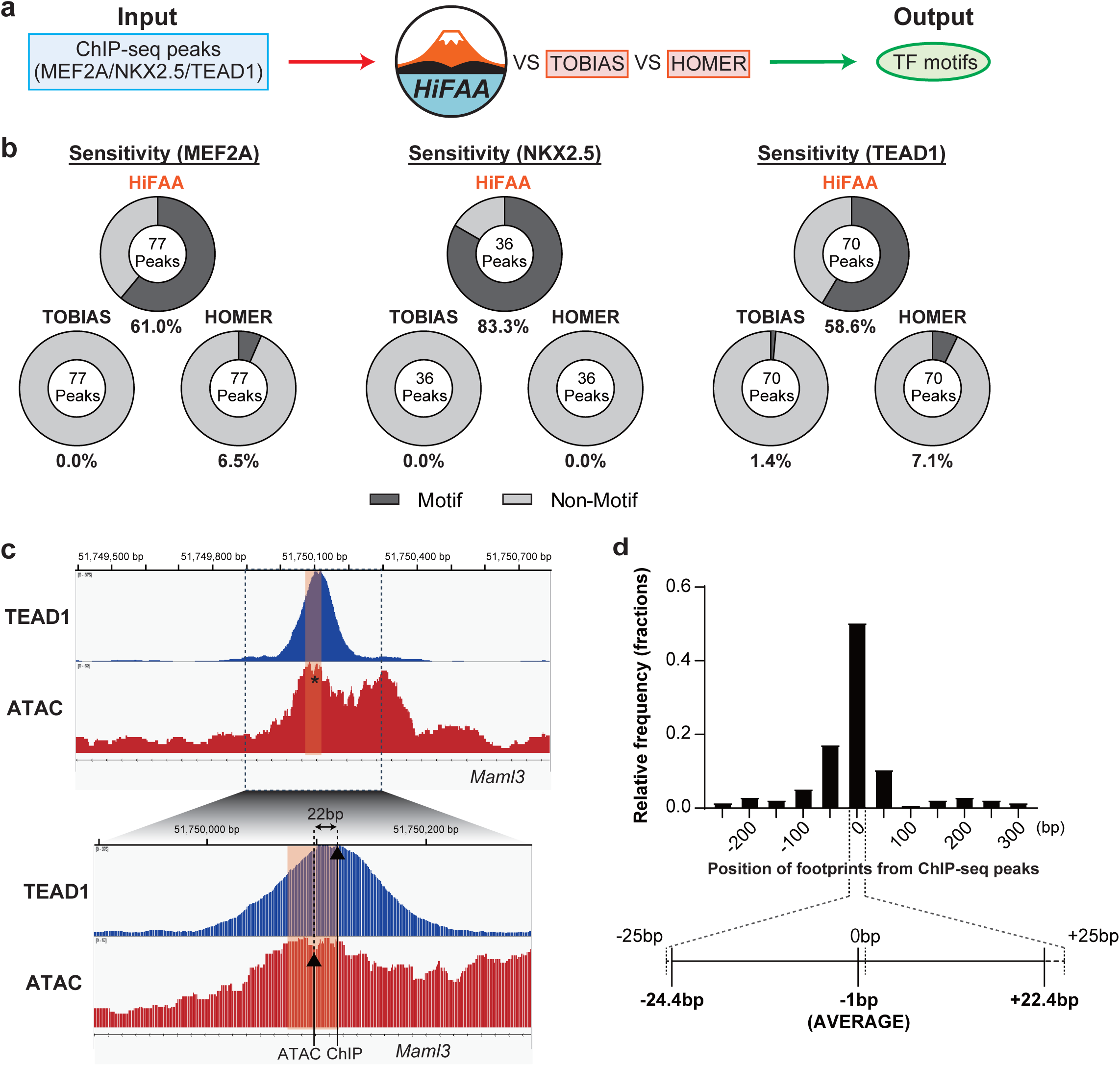
Sensitivity and spatial precision of motif detection by HiFAA, TOBIAS, and HOMER. **a,** Experimental scheme for evaluating motif detection sensitivity across MEF2A, NKX2.5, and TEAD1 in adult mouse heart. Publicly available ChIP-seq datasets (ChIP-seq: public datasets) were used to define reference TF-binding peaks, while ATAC-seq data were generated in this study (ATAC-seq: generated in this study) under matched conditions. Motif detection was performed using HiFAA, TOBIAS, and HOMER for direct comparison. **b,** Motif detection sensitivity summarized as donut charts, with dark gray indicating motif-containing peaks and light gray indicating motif absence. **c,** Example of TEAD1 footprint aligned with ChIP-seq peak (mm10, chr3:51,749,397–51,750,795). The footprint contains a TEAD1 motif and overlaps the ChIP-seq summit. **d,** Probability density distribution of distances between footprint bottoms and ChIP-seq summits across all peaks. The median deviation was ∼1 bp, with summits centered within the 47-bp HiFAA motif window, demonstrating near-nucleotide concordance.

HiFAA demonstrated striking improvements over existing methods (Fig. 2b). For MEF2A, HiFAA detected motifs in 47 of 77 peaks (61.0%), whereas HOMER detected motifs in only five peaks (6.5%) and TOBIAS detected none. For NKX2.5, HiFAA identified motifs in 30 of 36 peaks (83.3%), while HOMER and TOBIAS failed to detect any motifs. For TEAD1, HiFAA detected motifs in 41 of 70 peaks (58.6%), compared with five peaks (7.1%) for HOMER and one peak (1.4%) for TOBIAS. Thus, high-resolution footprints markedly improve detection sensitivity.

We further examined positional concordance between footprints detected by HiFAA Step 2 and ChIP-seq peaks (Fig. 2c). On average, the deviation between footprint bottoms and ChIP-seq peak summits was only 1 bp, and peak summits were consistently located near the footprint center. Accordingly, the HiFAA motif analysis window, defined as 47 bp centered on the footprint bottom, captured the vast majority of ChIP-seq peaks (Fig. 2d). Representative examples for MEF2A, NKX2.5, and TEAD1 showed ChIP-seq peaks located within HiFAA footprints (Extended data Fig. 1a). Motif enrichment analysis confirmed that the true motifs for each TF were ranked within the top five across replicates (MEF2A: 3rd, NKX2.5: 2nd, TEAD1: 5th; Extended data Fig. 1b). Thus, HiFAA Step 2 not only enhances sensitivity but also provides accurate statistical prioritization of biologically relevant motifs.

### Step 3: Footprint bottom height as a discriminator

Although HiFAA Step 2 improved sensitivity, it also identified motif-containing footprints lacking ChIP-seq support, raising concerns about specificity (Fig. 3a). To address this issue, we sought to identify structural parameters of footprints that could distinguish true TF-binding events (motif-positive footprints overlapping ChIP-seq peaks) from false positives (motif-positive footprints without ChIP-seq support). We evaluated four parameters: footprint depth, footprint width, footprint area, and footprint bottom height.

**Figure 3.**
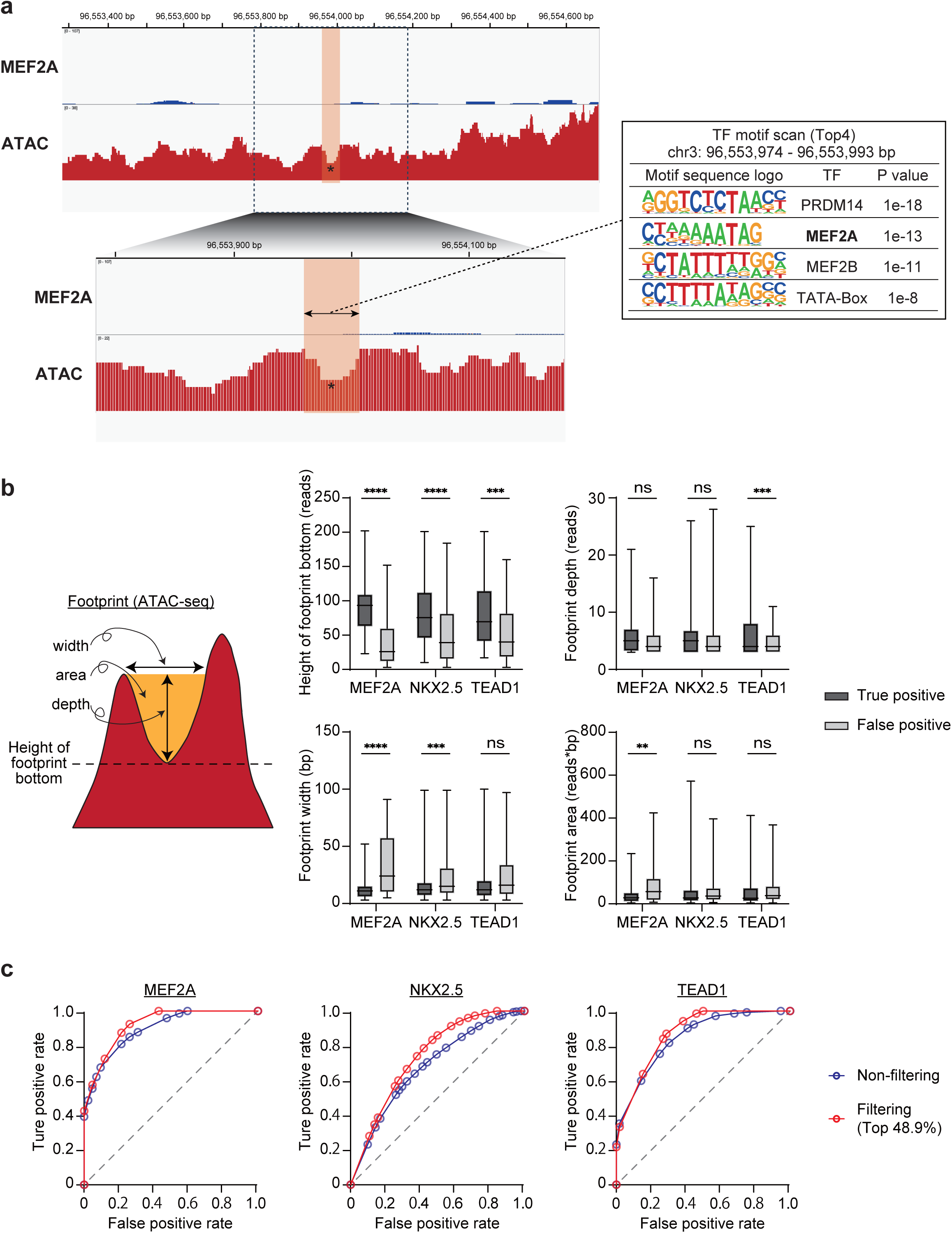
Evaluation of footprint metrics for distinguishing true from false positives. **a,** Example of a MEF2A motif-containing footprint lacking corresponding ChIP-seq support, illustrating a false-positive footprint. **b,** Comparison of four footprint metrics—bottom height, depth, width, and area—between true positives (motif+ and ChIP-seq+) and false positives (motif+ only) across MEF2A (n=44 vs 77), NKX2.5 (n=112 vs 341), and TEAD1 (n=56 vs 155). Only footprint bottom height consistently distinguished true from false footprints. Boxplots show interquartile range (IQR), median, and whiskers (1.5 × IQR). Significance was calculated by one-way ANOVA. Significance levels: **** P < 0.0001; *** P < 0.001; ** P < 0.01; * P < 0.05; ns, not significant. **c,** Receiver operating characteristic (ROC) curves evaluating footprint bottom height as a classifier. Filtering at the top 48.9% improved area under the curve (AUC) and sensitivity without compromising specificity across all transcription factors.

Using MEF2A, NKX2.5, and TEAD1 datasets, we found that only footprint bottom height consistently differentiated true from false positives across all TFs (Fig. 3b, Extended data Fig. 2a,b). Specifically, footprint bottoms were significantly higher in true binding sites than in false positives. By contrast, footprint depth, area, and width showed no significant differences between true and false positives. These results indicate that footprint bottom height serves as a robust index for identifying genuine TF-binding events.

### Step 3: Ranking and selection of high-confidence footprints

Building on this observation, we introduced footprint ranking and filtering based on bottom height in HiFAA Step 3. We evaluated performance using receiver operating characteristic (ROC) analysis, defining true positives as motif-containing footprints overlapping ChIP-seq peaks and false positives as motif-containing footprints lacking ChIP-seq support. Thresholds were varied by percentile ranking of footprint bottom height within each TAD.

This analysis demonstrated that filtering footprints above the 48.9th percentile optimized performance across TFs. For MEF2A, the area under the ROC curve (AUC) increased from 0.8962 (no filtering) to 0.9211 with filtering, while sensitivity increased to 87.5% without compromising specificity (Fig. 3c, Supplementary Data 1). For NKX2.5, filtering improved the AUC from 0.6814 to 0.7388, with modest gains in sensitivity and specificity (Fig. 3c, Supplementary Data 2). For TEAD1, filtering increased the AUC from 0.8470 to 0.8718 and sensitivity from 75.4% to 84.1%, while specificity was maintained (Fig. 3c, Supplementary Data 3). These results strongly support the effectiveness of footprint bottom height as a discriminative metric and validate footprint ranking and selection as a critical step in HiFAA.

### Genome-scale evaluation in the heart

To assess HiFAA’s practical utility, we compared its performance against HOMER and TOBIAS using ATAC-seq and ChIP-seq data from adult mouse hearts^16^ (Fig. 4a). We first identified the top 100 human heart-enriched genes from GTEx RNA-seq data. For validation, publicly available ChIP-seq datasets for MEF2A, NKX2.5, and TEAD1 were obtained and processed to define reference TF-binding peaks. In parallel, we generated independent ATAC-seq datasets from adult mouse hearts under matched conditions. These ATAC-seq data were analyzed with HiFAA, HOMER, and TOBIAS, and predicted TF-binding regions were compared with ChIP-seq peaks to calculate sensitivity and specificity.

**Figure 4.**
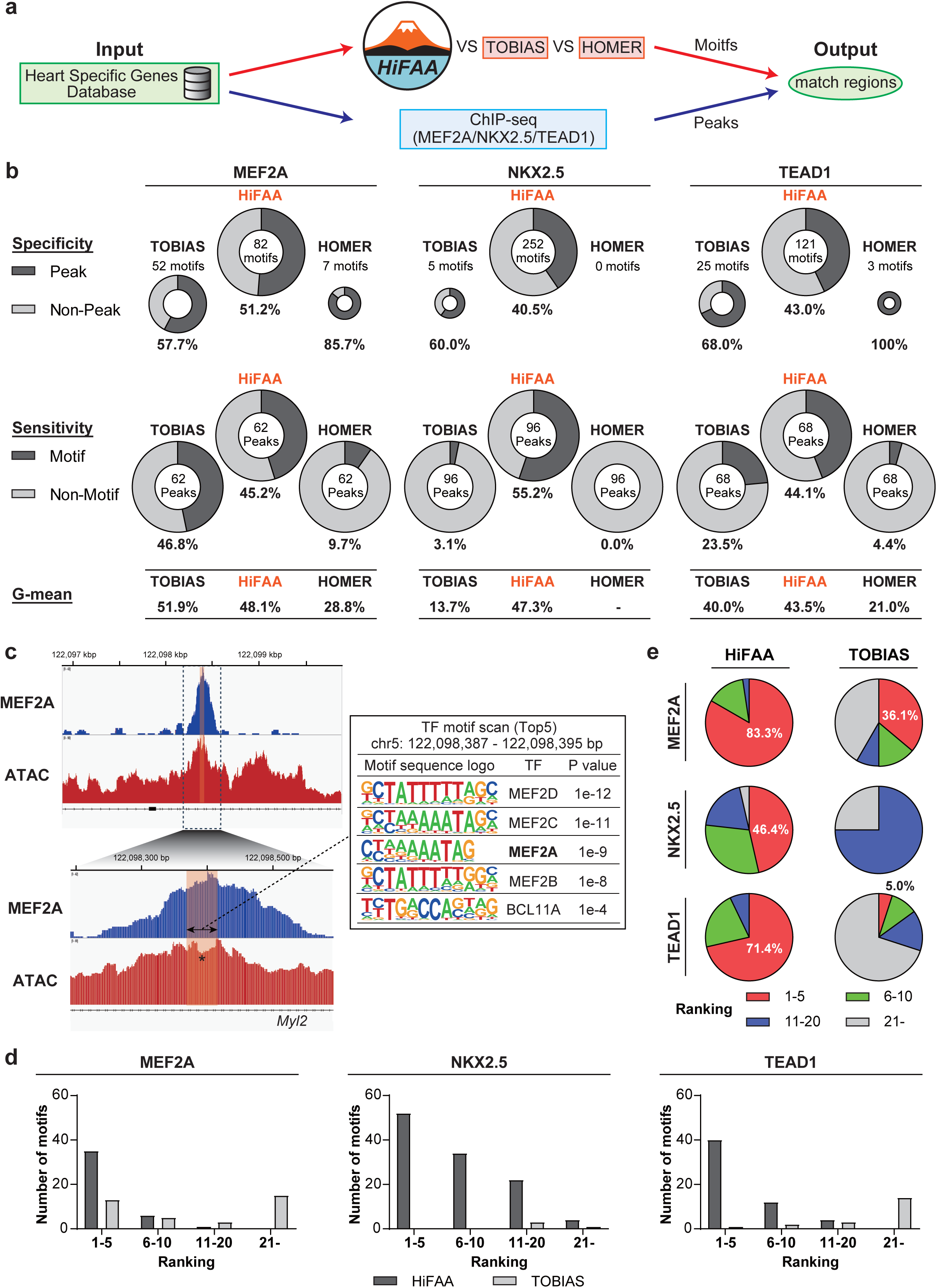
Comparative evaluation of HiFAA, TOBIAS, and HOMER using heart-enriched gene sets. **a,** Schematic overview of motif detection performance evaluation using HiFAA, TOBIAS and HOMER, integrating heart-specific gene information. A set of 100 heart-specific genes was selected from GTEx-derived human heart RNA-seq data. Publicly available ChIP-seq datasets (ChIP-seq: public datasets) for MEF2A, NKX2.5, and TEAD1 were used to define reference TF binding peaks, while ATAC-seq data were generated in this study (ATAC-seq: generated in this study) under matched conditions. Overlaps between motif-positive footprints (from HiFAA, TOBIAS or HOMER) and ChIP-seq peaks within the heart-specific gene set were assessed to compute sensitivity and specificity for each method. **b,** Donut charts summarizing sensitivity and specificity for each method. HiFAA showed consistently higher sensitivity while maintaining stable specificity. Geometric-mean values are shown to reflect overall classification performance, where HiFAA achieved consistently higher and more stable scores across TFs. **c,** Representative browser view of MEF2A footprint overlapping a ChIP-seq summit. **d,** Distribution of motif ranking positions across transcription factors, showing enrichment of HiFAA motifs in the top 1– 5 ranks. **e,** Proportion of motif rankings by HiFAA and TOBIAS. HiFAA consistently prioritized true motifs in higher positions.

HOMER achieved high specificity across TFs but exhibited very low sensitivity (sensitivity 0–9.7%, specificity 85.7–100%; Fig. 4b). TOBIAS showed modest improvements in sensitivity and stable specificity (sensitivity 3.1–46.8%, specificity 57.7–68.0%). In contrast, HiFAA consistently demonstrated higher sensitivity (44.1–55.2%) with stable specificity (40.5–51.2%). Importantly, when combining both sensitivity and specificity into a single balanced metric (geometric mean, G-mean), HiFAA consistently achieved the highest G-mean values across TFs (43–48%), whereas HOMER showed low values due to poor sensitivity and TOBIAS showed unstable values. This demonstrates HiFAA’s robustness in balancing sensitivity and specificity.

Representative examples confirmed that HiFAA-predicted footprints coincided with ChIP-seq peaks, demonstrating that HiFAA footprints reflect genuine TF binding rather than mere motif presence (Fig. 4c, Extended data Fig. 6a,b). Motif enrichment analysis further revealed that HiFAA ranked target motifs within the top five in the majority of footprint regions (46.4–83.3%), whereas TOBIAS frequently ranked target motifs below the top ten (0–36.1%) (Fig. 4d, e). This highlights HiFAA’s ability not only to detect correct motifs but also to prioritize them appropriately, thereby streamlining downstream functional analyses.

Further, we present a representative example demonstrating the effectiveness of the footprint bottom height filtering introduced in HiFAA step 3 (Extended data Fig. 7). In this example, a region with a footprint-like shape in ATAC-seq and a target TF motif was detected by motif analysis, but no actual ChIP-seq peak was present. While this region would have been considered positive by conventional methods and HiFAA Step 2, it was excluded by HiFAA Step 3 because its footprint bottom height was in the top 49.7% of TADs, which was below the threshold. Thus, HiFAA Step 3 effectively eliminated false-positive regions with poor biological basis despite being motif-positive.

### Detection of developmental stage–specific TF binding

Finally, we evaluated whether HiFAA could capture dynamic TF-binding events across species, tissues, and developmental stages. We focused on the fetal-to-adult hemoglobin switch in human erythropoiesis, a process regulated by the TF BCL11A. In adult hematopoietic cells, BCL11A represses fetal β-hemoglobin expression, and its own expression is induced by GATA1 binding to an enhancer element^12,13^.

We analyzed publicly available ATAC-seq and GATA1 ChIP-seq datasets from erythroid cells derived from fetal liver CD34⁺ hematopoietic progenitors and from adult peripheral blood^17^. In fetal samples, HiFAA identified two motif-positive footprints, both overlapping ChIP-seq peaks (specificity 100%, sensitivity 50.0%; G-mean 70.7%), whereas HOMER and TOBIAS detected none (Fig. 5a). In adult samples, HiFAA detected six motif-positive footprints, all overlapping ChIP-seq peaks (specificity 100%, sensitivity 42.9%; G-mean 65.5%). HOMER again detected no motifs, and TOBIAS detected only two motif-positive footprints (specificity 100%, sensitivity 14.3%; G-mean 37.8%). These results demonstrate that HiFAA maintains stable G-mean values across biological conditions, highlighting its ability to balance sensitivity and specificity while capturing developmental changes in TF binding.

**Figure 5.**
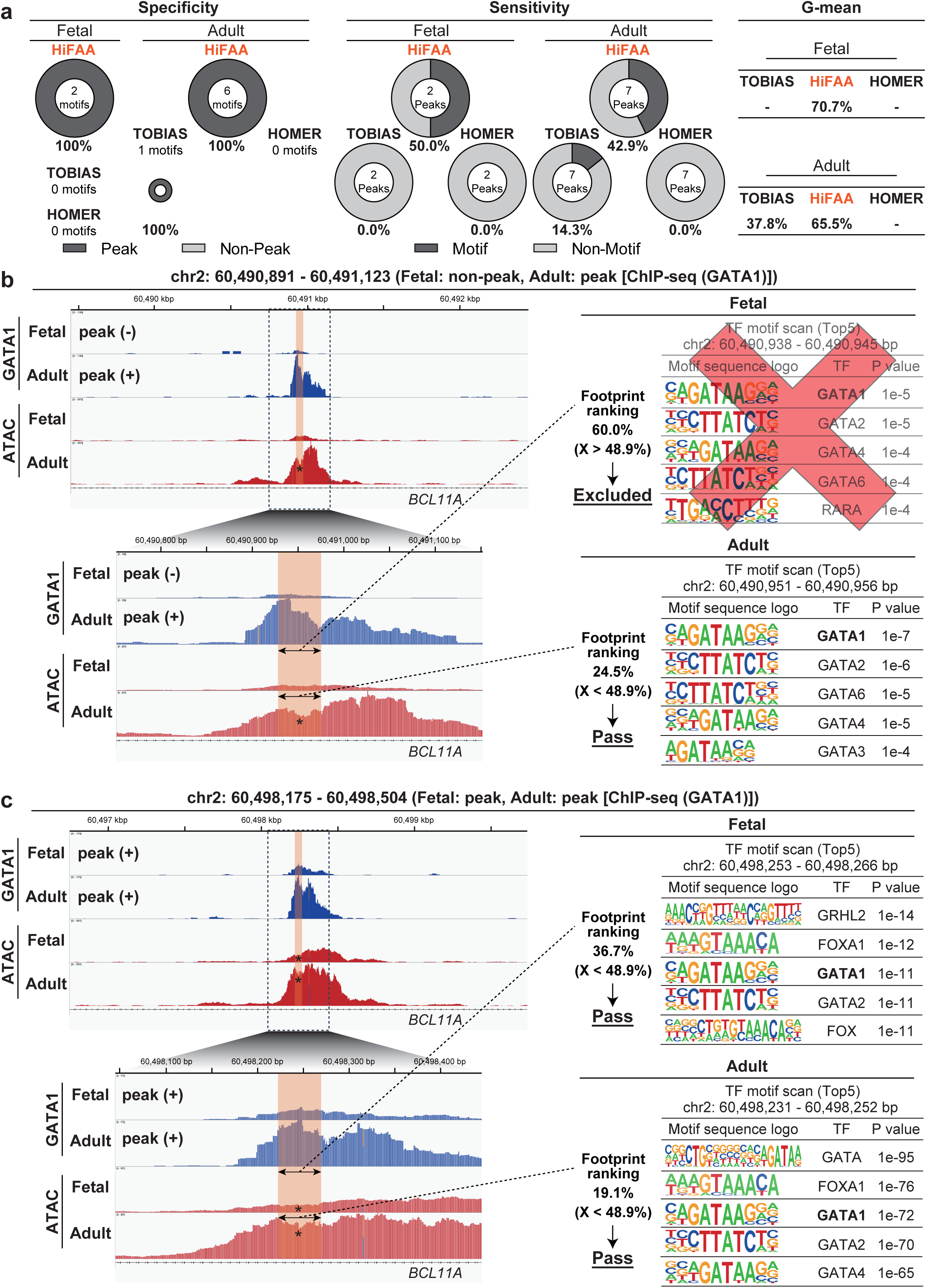
Developmental-stage-specific detection of GATA1 footprints. **a,** Motif detection specificity and sensitivity for GATA1 in fetal and adult erythroid progenitors visualized as donut charts. HiFAA consistently detected motif-positive footprints overlapping ChIP-seq peaks, whereas HOMER and TOBIAS performed poorly. HiFAA also achieved the highest Geometric-mean scores across both conditions, indicating balanced sensitivity and specificity. **b,** Example of a stage-specific GATA1 footprint at the BCL11A enhancer. A footprint was detected and ranked within the top 24.5% in adult cells but excluded in fetal cells due to lower ranking (60.0%). **c,** Example of a conserved GATA1 footprint detected in both fetal and adult cells, ranked within the top 48.9% in both cases. These results illustrate HiFAA’s capacity to capture both dynamic and stable TF binding across development.

HiFAA also successfully detected stage-specific changes in GATA1 binding. For example, in one enhancer region, no GATA1 ChIP-seq peak was present in fetal samples, but a clear peak appeared in adults (Fig. 5b). HiFAA identified a motif-positive footprint in adults (ranked top 24.5% of footprints) but excluded the corresponding fetal footprint due to low reliability (ranked top 60.0%), thereby accurately capturing the developmental switch. In regions with consistent GATA1 binding across stages, HiFAA identified motif-positive footprints in both fetal and adult samples (top 36.7% and 19.1%, respectively; Fig. 5c). These findings demonstrate that HiFAA reliably detects both stable and dynamic TF-binding events across species and developmental stages.

## Discussion

In this study, we developed HiFAA, a computational framework designed to improve the accuracy of TF-binding inference from ATAC-seq data. HiFAA introduces three methodological innovations: TAD-restricted analysis, high-resolution footprints, and ranking by footprint bottom height. Together, these steps increased both sensitivity and geometric mean (G-mean) compared with HOMER^6^ and TOBIAS^7^. Unlike HiFAA, HOMER identifies motifs by scanning peak centers—computationally efficient but insensitive. TOBIAS introduced footprint valleys, improving sensitivity but relying on width/depth metrics. HiFAA builds on these foundations while integrating chromatin context, single-base resolution, and ranking to overcome limitations. Benchmarking against ChIP-seq data^16^ for MEF2A, NKX2.5, and TEAD1 in mouse cardiac tissue demonstrated that HiFAA detected true TF-binding motifs with significantly higher sensitivity than either HOMER or TOBIAS. Importantly, HiFAA footprints showed near-perfect positional concordance with ChIP-seq peaks, with footprint bottoms aligning closely to ChIP-seq summits. Ranking footprints by bottom height provided a robust criterion for distinguishing true binding sites from false positives, further improving specificity. To more rigorously evaluate overall performance, we also calculated G-mean of sensitivity and specificity. HiFAA consistently achieved high and stable G-mean values across transcription factors and developmental contexts, indicating that it maintains an optimal balance between sensitivity and specificity. In contrast, HOMER showed low G-mean values due to extremely poor sensitivity despite high specificity, while TOBIAS achieved higher G-mean in some cases (e.g., MEF2A) but unstable and generally lower values for other TFs. This highlights HiFAA’s advantage as a method that avoids trading sensitivity for specificity, achieving balanced discrimination. Finally, HiFAA captured dynamic TF binding changes during development, such as stage-specific GATA1 binding at the BCL11A enhancer during the fetal-to-adult hemoglobin switch^12,13^, highlighting its ability to resolve fine-grained regulatory dynamics across species and developmental stages.

The superior performance of HiFAA relative to HOMER and TOBIAS can be attributed to the integration of the methodological innovations. HOMER’s strength lies in its computational simplicity, robustness, and high specificity, making it a reliable first-line tool for motif enrichment analyses in many studies^6^. However, its reliance on peak-centered scanning often results in low sensitivity because motifs in open chromatin may not correspond to true binding events. TOBIAS, on the other hand, improved sensitivity by explicitly modeling footprint valleys and has been successfully applied to genome-wide analyses and developmental contexts^7^. Nonetheless, its metrics based mainly on footprint depth and width can misclassify local chromatin variation as binding. In contrast, HiFAA capitalizes on higher-order chromatin architecture (via TAD restriction), enhanced structural resolution (via single-base footprints), and a biologically meaningful ranking system (via bottom height). This combination reduces background noise, improves signal detection, and prioritizes true binding sites. The consistently high G-mean values observed for HiFAA provide quantitative evidence of this balanced performance, complementing ROC/AUC metrics by directly capturing threshold-dependent trade-offs.

Restricting the analysis to the TAD containing the target gene provides both biological and computational advantages. From a computational standpoint, TAD restriction dramatically reduces the size of the genomic search space. The average size of a TAD in mammals ranges from 200 kb to 2 Mb^9,10^, representing only a small fraction of the genome. Thus, restricting analysis to a single TAD can reduce the sequence space by more than 90% relative to genome-wide scanning. From a biological perspective, enhancer–promoter interactions are predominantly confined within TADs. Previous studies suggest that more than 80–90% of TF-binding sites relevant to gene regulation occur within the same TAD as the target gene^3,4,9^. By focusing analysis within this domain, HiFAA ensures that maximal functionally relevant regulatory interactions are captured while discarding distal binding sites that are less likely to influence the gene of interest. Although this could potentially miss long-range inter-TAD interactions, the trade-off strongly favors specificity and interpretability.

Step 2 of HiFAA introduces single-base-resolution footprints derived from BAM data and pseudo-reads. This innovation enables more accurate delineation of footprint structure compared with existing methods, which define footprints primarily by width and depth^7,18^. Our analyses revealed that footprint bottoms aligned almost perfectly with ChIP-seq peak summits, with an average deviation of only 1 bp. This remarkable concordance underscores the biological validity of HiFAA’s high-resolution footprints and demonstrates that TF-binding events leave precise and detectable signatures in ATAC-seq data^5^.

HiFAA’s third innovation, ranking footprints based on bottom height, provides a robust mechanism to discriminate true binding sites from false positives. The biological plausibility of this metric lies in the distinction between footprints within open chromatin and regions of closed chromatin. When the footprint bottom is extremely low, the apparent depletion of Tn5 insertions is more likely due to globally inaccessible chromatin rather than protection by a bound TF. In contrast, genuine TF footprints appear as local depressions embedded within highly accessible regions, where the footprint bottom remains at a relatively elevated level compared with surrounding closed chromatin. Thus, a footprint with a sufficiently high bottom within an ATAC-seq peak is more indicative of TF occupancy than one that simply reflects overall chromatin compaction. Our analyses confirmed that such high-bottom footprints were more likely to overlap with ChIP-seq peaks, underscoring that bottom height serves as a biologically meaningful indicator of true TF binding. By leveraging this principle, HiFAA’s ranking procedure effectively converts a raw structural parameter into a reliable criterion for prioritizing high-confidence binding sites. Importantly, this approach not only improves individual metrics but, as reflected in the G-mean, ensures that sensitivity gains do not come at the cost of specificity.

Our analysis of GATA1 binding at the BCL11A enhancer during the fetal-to-adult hemoglobin switch exemplifies this capability. HiFAA not only detected stable GATA1 binding across developmental stages but also identified stage-specific footprints corresponding to adult-specific enhancer activation, consistent with prior work on the regulatory role of BCL11A in hemoglobin switching^12,13^. This ability to resolve dynamic TF binding has broad implications, enabling HiFAA to be applied to diverse biological contexts such as lineage specification^19^ and disease-associated regulatory reprogramming^20^.

Despite its advantages, HiFAA has several limitations. First, by restricting analysis to TADs, HiFAA may miss distal regulatory interactions across TAD boundaries^9,10^. Second, HiFAA is limited to known motifs and cannot discover novel motifs de novo^21^. Third, HiFAA relies on high- quality ATAC-seq data; shallow sequencing reduces footprint resolution, which may compromise sensitivity^5^.

Future work will aim to address these limitations. Integration of de novo motif discovery^21^ into HiFAA would extend its scope beyond known TFs. Incorporation of three-dimensional chromatin architecture data, such as Hi-C or Micro-C^22^, could capture long-range enhancer– promoter interactions. Furthermore, adaptation of HiFAA to single-cell ATAC-seq^23^ would allow dissection of TF-binding heterogeneity at cellular resolution.

In conclusion, HiFAA represents a significant advance in ATAC-seq–based motif analysis by integrating biologically informed restriction of search space, high-resolution footprinting, and robust ranking of footprints. Importantly, HiFAA complements rather than replaces prior tools: HOMER remains valuable for its computational efficiency and specificity, and TOBIAS provides genome-wide sensitivity and has been widely adopted in developmental studies. HiFAA extends these approaches by integrating their strengths while addressing their weaknesses, thereby achieving a more balanced trade-off between sensitivity and specificity. By consistently delivering high G-mean values, HiFAA demonstrates that it not only improves raw sensitivity but also maintains the delicate balance with specificity, setting a new standard for robust TF-binding inference. Beyond static motif detection, HiFAA enables the discovery of dynamic, developmentally regulated TF-binding events, offering new insights into the transcriptional regulatory logic of cellular differentiation. HiFAA establishes a new standard for footprint analysis and opens promising avenues for advancing our understanding of gene regulation in health and disease.

## Acknowledgments

This work was supported by grants from the Japan Society for the Promotion of Science KAKENHI [20K17151, 22K08204, 23K24327, 25K11386], grants from the Japan Agency for Medical Research and Development (AMED) [23bm1123026h0001], and by the Suzuki Kenzo Memorial Foundation for the Advancement of Medical Sciences, the Osaka Medical Research Foundation for Intractable Diseases, SENSHIN Medical Research Foundation, Japan Heart Foundation Research Grant on Dilated Cardiomyopathy, The NOVARTIS Foundation (Japan) for the Promotion of Science, Miyata Foundation Bounty for Pediatric Cardinovasucular Research, Mochida Memorial Foundation for medical and Pharmaceutical Research, and Kobayashi Foundation.

## Author contributions

H.I. and K.M. conceptualized the project and designed the experiments. H.I. developed the HiFAA software. C.O. and H.K. performed additional bioinformatics analysis. H.I., O.T. and K.M. performed ATAC-seq experiments. H.I., S.T., and K.M. engaged in extensive discussions regarding the structure and framing of the manuscript. S.T. and K.M. analyzed the data and wrote the manuscript with inputs from all authors. S.T., and K.M. secured funding and coordinated the collaborative efforts.

