## Supplementary Information for "HiFAA: a high-confidence framework for transcription factor footprinting with ATAC-seq"

### Extended Data Fig. 1

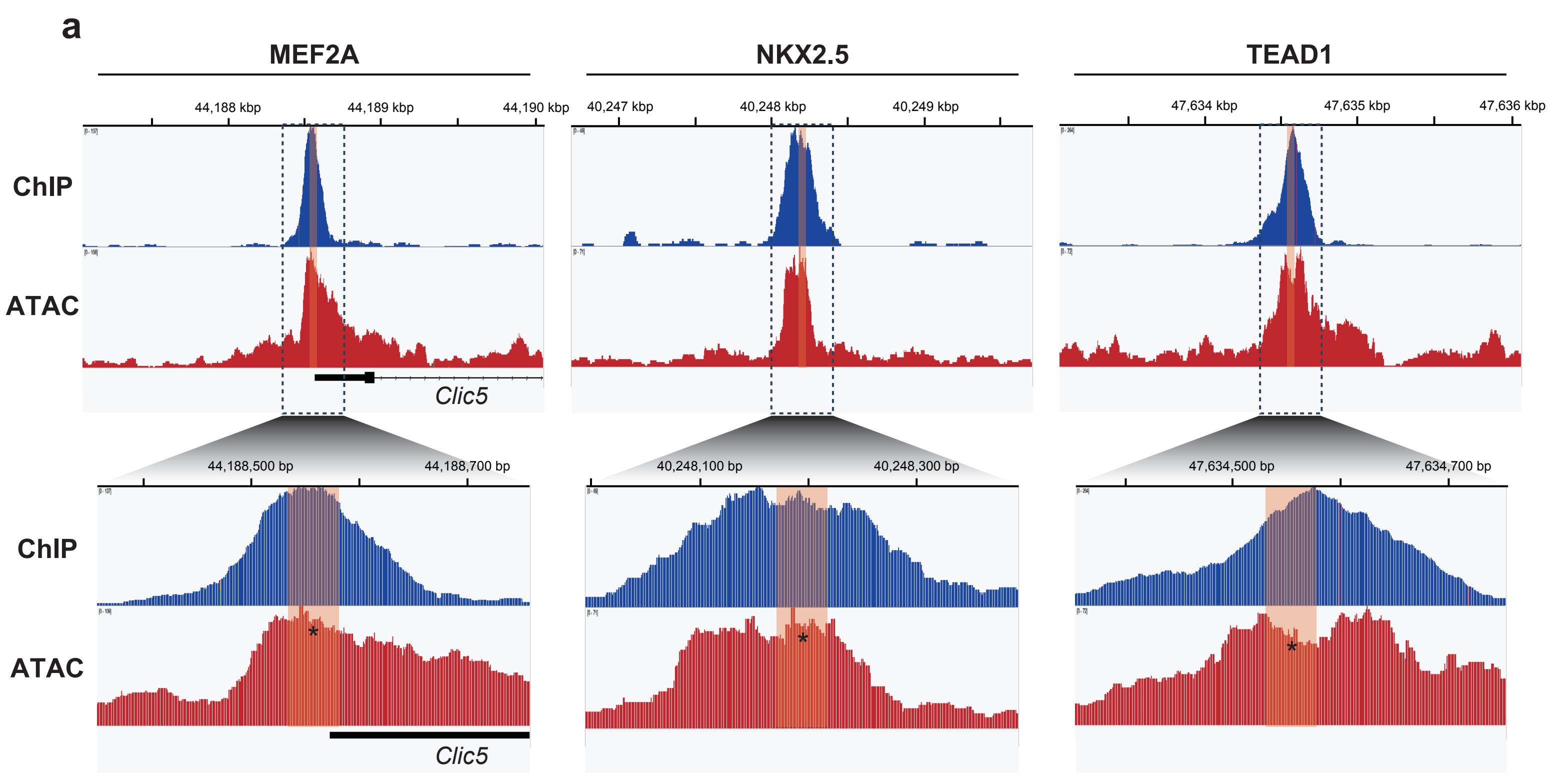


**b**

TF motif scan (Top5)
chr17: 44,188,551 - 44,188,561 bp

| Motif sequence logo | TF | P value |
|---|---|---|
| 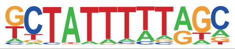 | MEF2D | 1e-15 |
| 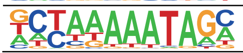 | MEF2C | 1e-14 |
| 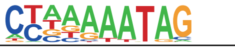 | **MEF2A** | 1e-13 |
| 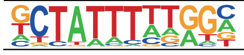 | MEF2D | 1e-12 |
| 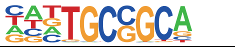 | ZFP57 | 1e-5 |

TF motif scan (Top5)
chr12: 40,248,189 - 40,248,194 bp

| Motif sequence logo | TF | P value |
|---|---|---|
| 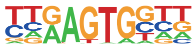 | BAPX1 | 1e-5 |
| 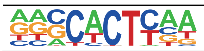 | **NKX2.5** | 1e-5 |
| 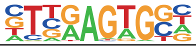 | NKX2.2 | 1e-5 |
| 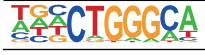 | ZNF416 | 1e-5 |
| 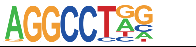 | ZFX | 1e-4 |

TF motif scan (Top5)
chr14: 47,634,551 - 47,634,555 bp

| Motif sequence logo | TF | P value |
|---|---|---|
| 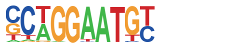 | TEAD2 | 1e-9 |
| 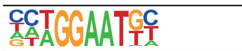 | TEAD | 1e-9 |
| 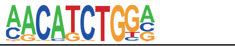 | ZBTB18 | 1e-9 |
| 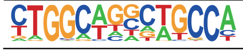 | TLX | 1e-8 |
| 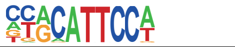 | **TEAD1** | 1e-8 |

**Extended Data Fig. 1. Footprints are spatially aligned with ChIP-seq summits. a,** Representative footprints for MEF2A, NKX2.5, and TEAD1 in adult mouse heart showing ATAC-seq signal valleys aligned with ChIP-seq summits. Orange bars denote 47-bp footprint windows. **b,** Motif enrichment analysis of footprint windows revealed that corresponding motifs were consistently ranked within the top five.

### Extended Data Fig. 2

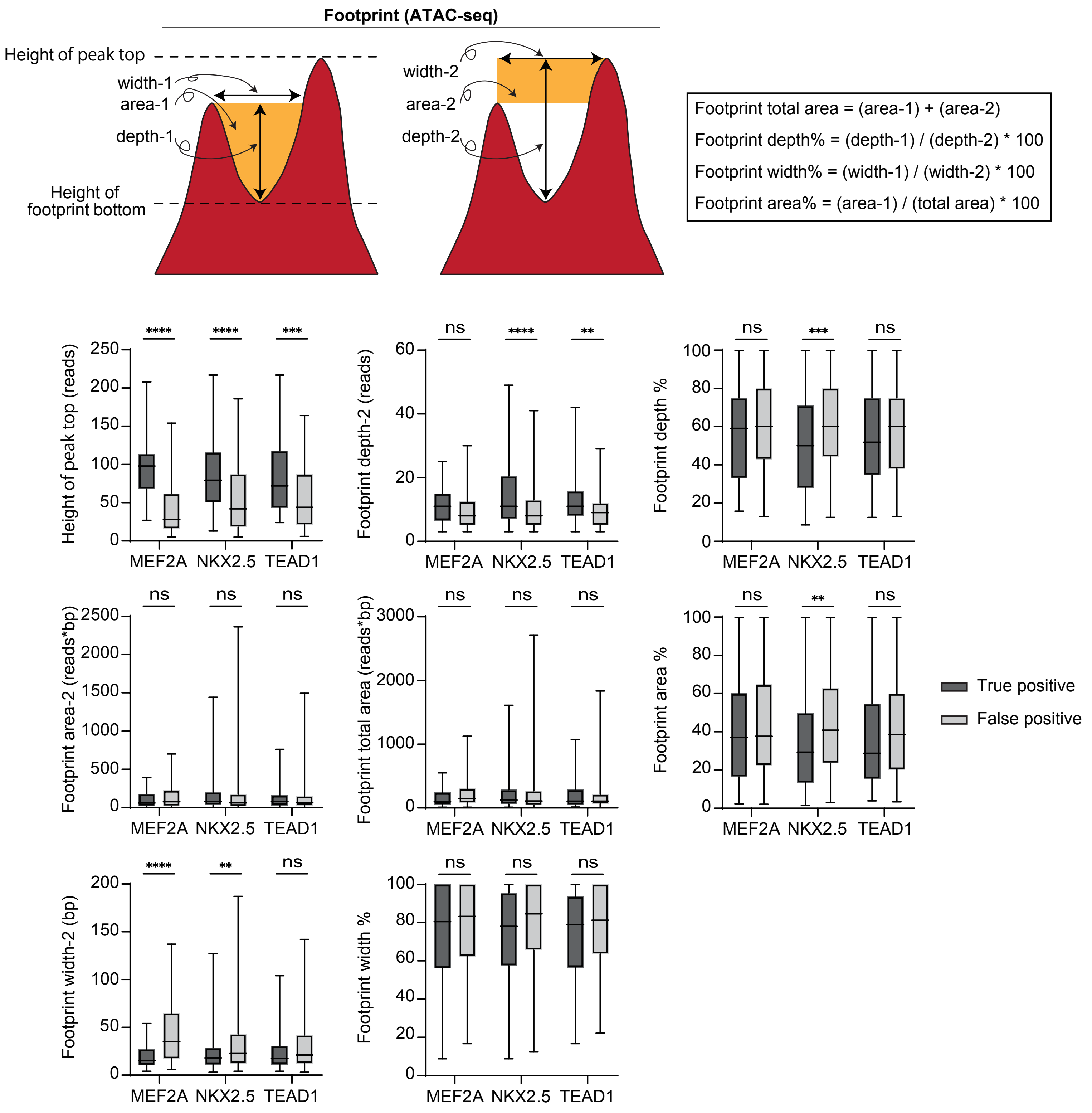


**Extended Data Fig. 2. Comparative analysis of footprint metrics.** Quantification of multiple footprint parameters between true positives (motif+ and ChIP-seq+) and false positives (motif+ only). Only footprint bottom height showed consistent discriminatory power across MEF2A, NKX2.5, and TEAD1.

### Extended Data Fig. 3

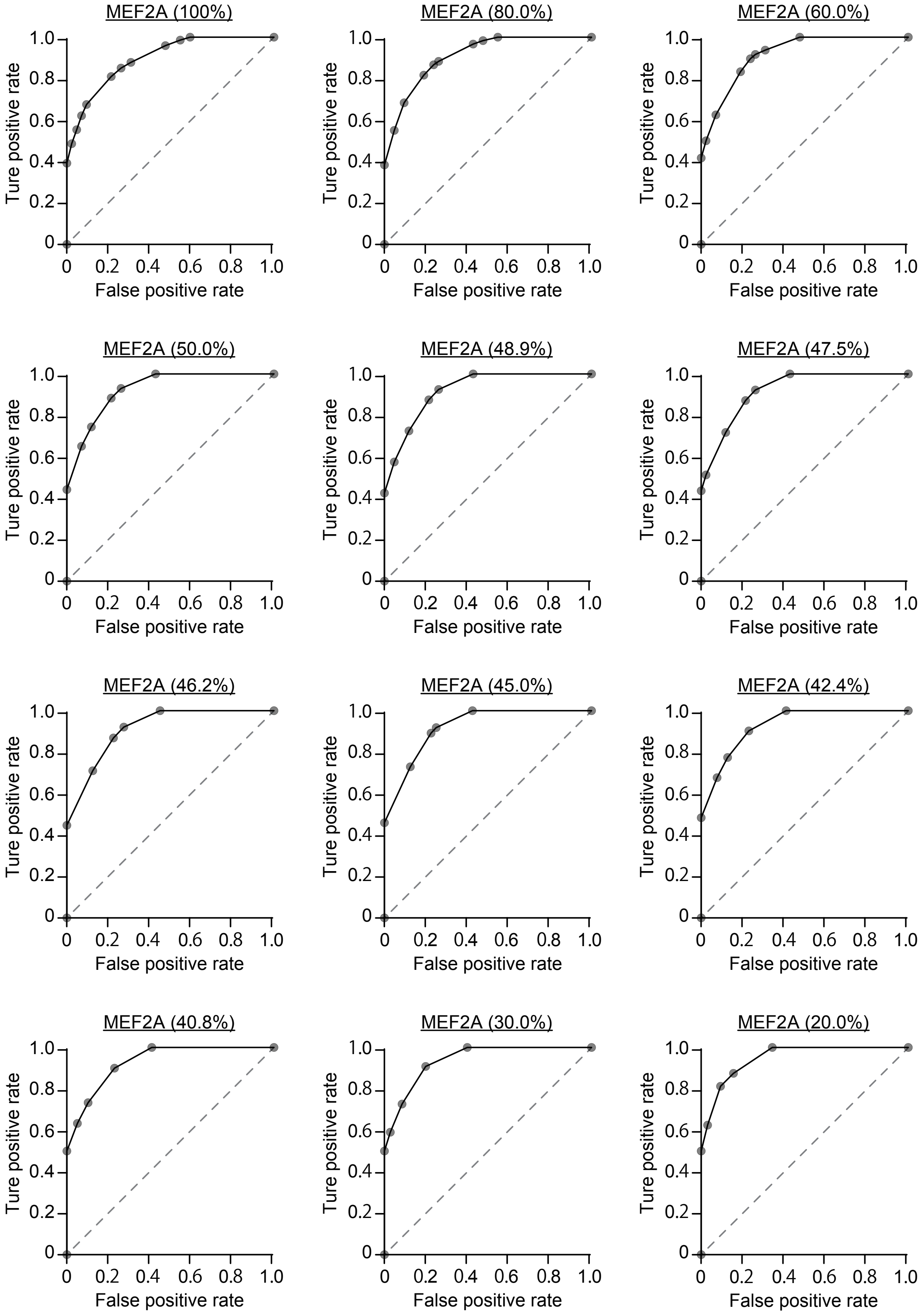


**Extended Data Fig. 3.** ROC analysis for MEF2A footprint classification. ROC curves evaluating footprint bottom height thresholds. The top 48.9% cutoff achieved the highest classification performance, balancing sensitivity and specificity.

### Extended Data Fig. 4

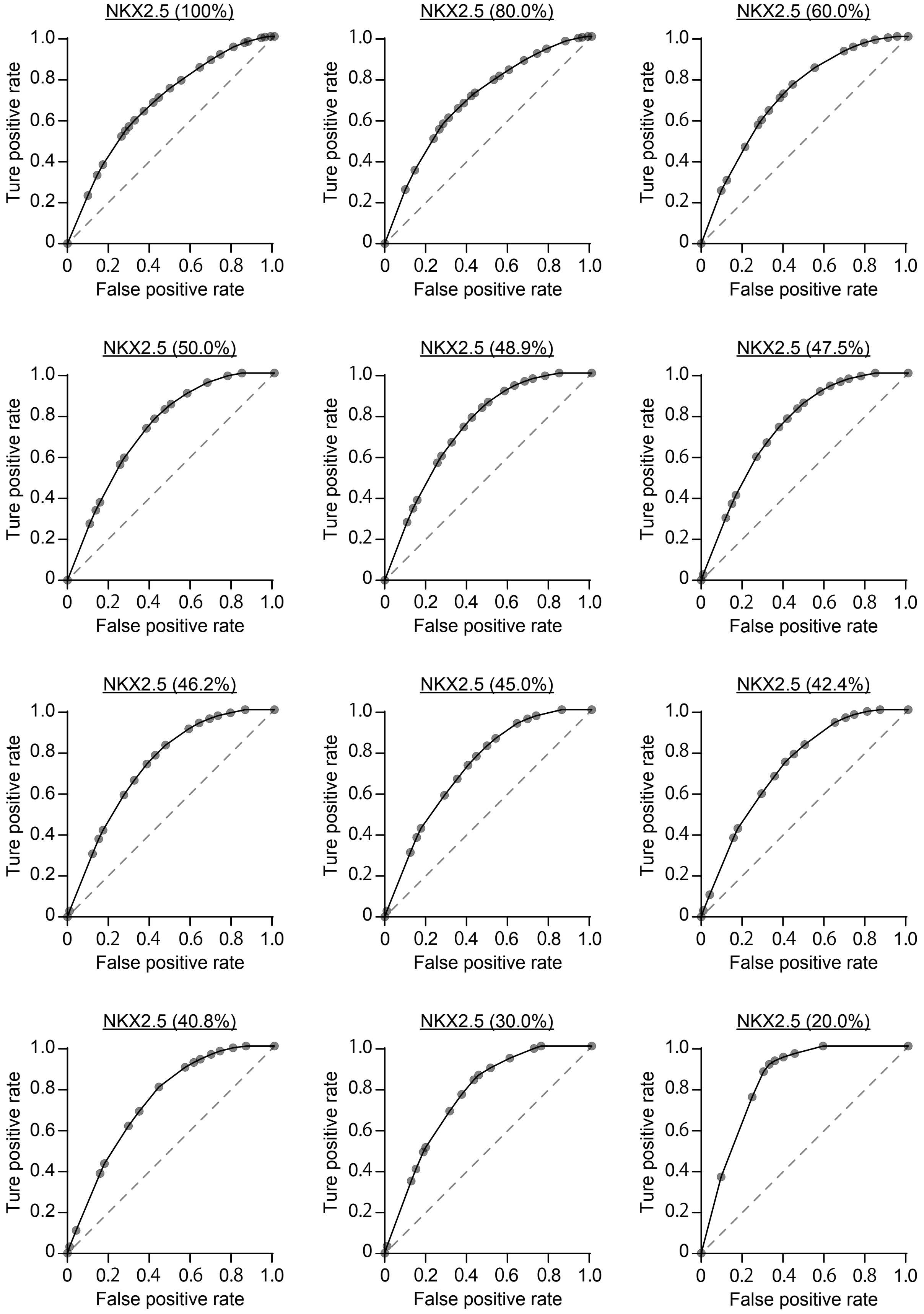


**Extended data Fig. 4.** ROC analysis for NKX2.5 footprint classification. ROC curves evaluating footprint bottom height thresholds. The optimal cutoff was the top 48.9% of footprints, improving AUC and reducing false positives.

### Extended Data Fig. 5

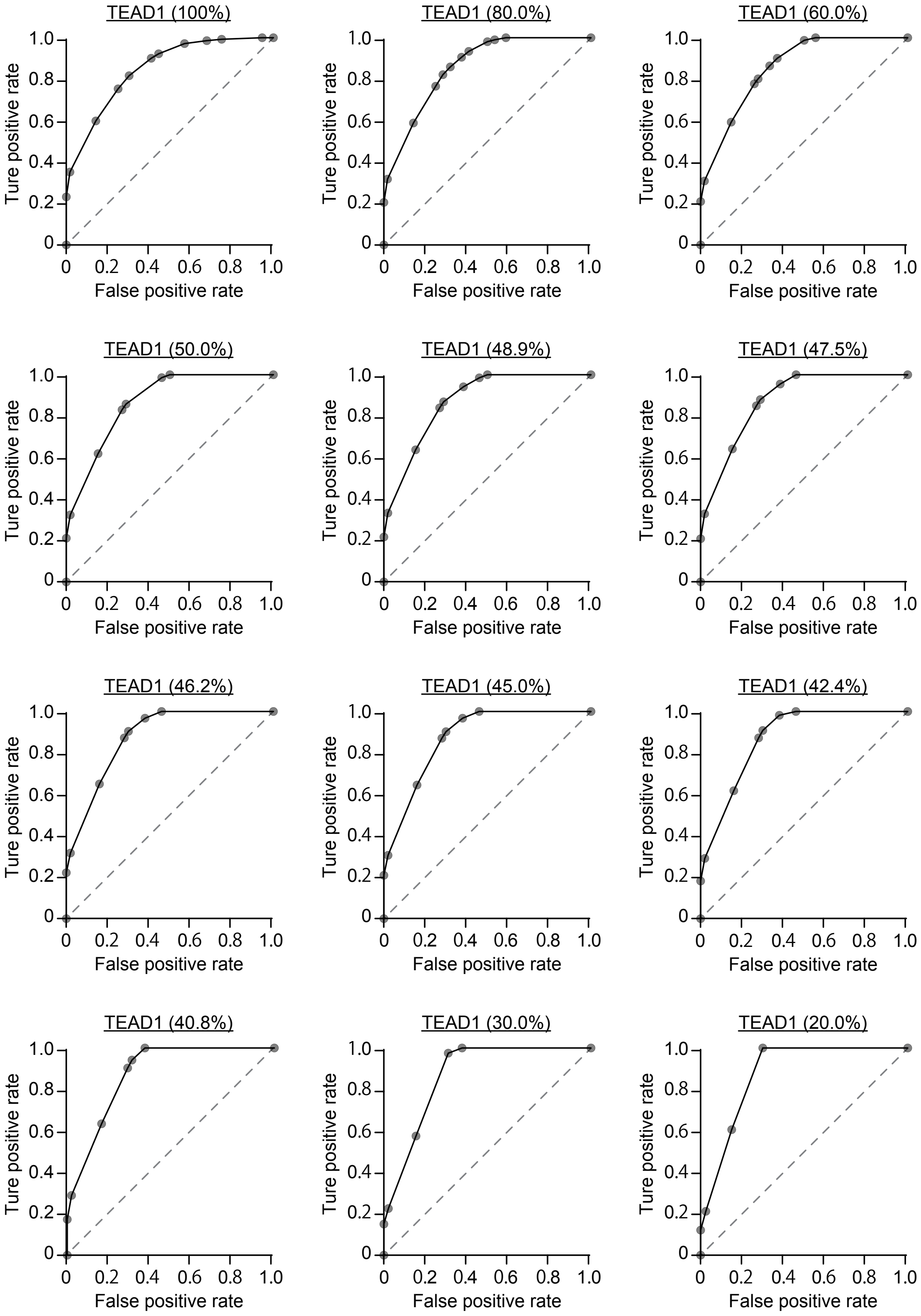


**Extended data Fig. 5**. ROC analysis for TEAD1 footprint classification. ROC curves for evaluating footprint bottom height thresholds. The 48.9% cutoff consistently improved classification accuracy.

### Extended Data Fig. 6

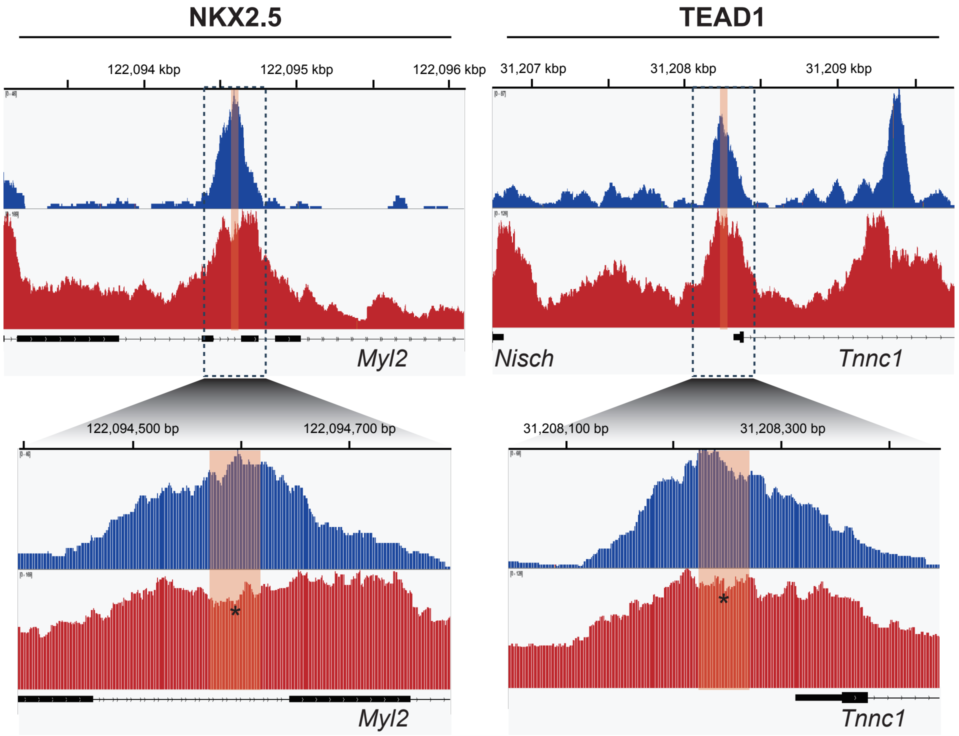


TF motif scan (Top5)
chr5: 122,094,594 - 122,094,596 bp

| Motif sequence logo | TF | P value |
|---|---|---|
| | EDF2 | 1e-4 |
| | BAPX1 | 1e-4 |
| | NKX3.1 | 1e-4 |
| | MYOG | 1e-4 |
| | **NKX2.5** | 1e-4 |

TF motif scan (Top5)
chr14: 31,208,245 - 31,208,249 bp

| Motif sequence logo | TF | P value |
|---|---|---|
| | ZFP809 | 1e-11 |
| | STAT3 | 1e-9 |
| | STAT3 | 1e-9 |
| | STAT4 | 1e-9 |
| | **TEAD1** | 1e-9 |

**Extended data Fig. 6. ATAC-seq footprints overlap with ChIP-seq peaks and recover corresponding motifs.** Representative examples for NKX2.5 and TEAD1 footprints aligned with ChIP-seq summits. Motif analysis ranked corresponding TF motifs among the top five hits.

### Extended Data Fig. 7

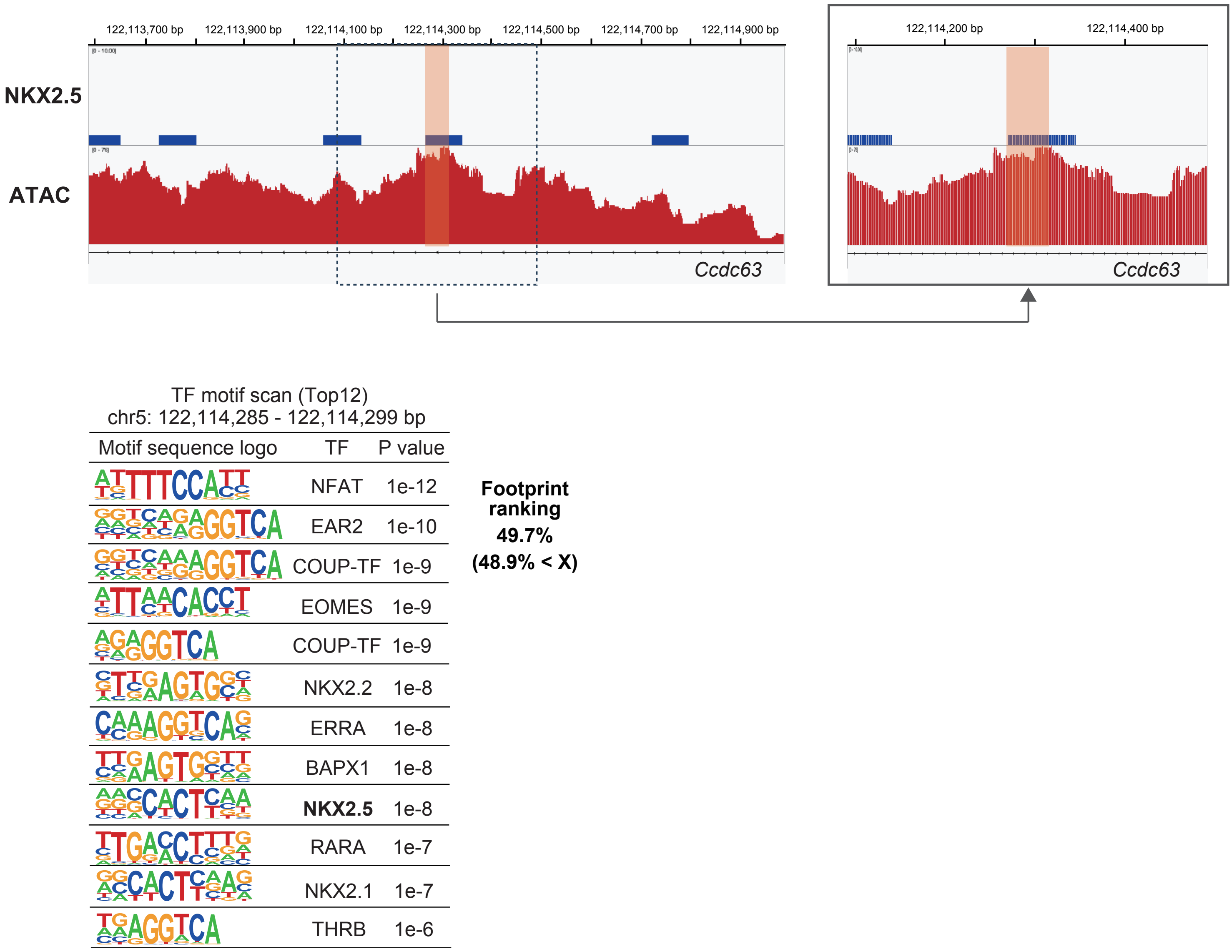


**Extended data Fig. 7. HiFAA excludes low-confidence footprints by bottom height filtering.** Example of an NKX2.5 motif-containing footprint lacking ChIP-seq support. Although motif-positive, the footprint ranked below the 48.9% cutoff and was excluded, demonstrating HiFAA's ability to filter low-confidence predictions.

Supplementary Table 1. Reagents and Antibodies

| Name | Product No. | Vendor |
|---|---|---|
| anti-PCM-1 antibody | HPA023370 | Sigma-Aldrich |
| Alexa Fluor 488-conjugated secondary antibody | A-11034 | Invitrogen |
| ATAC-Seq Kit | 53150 | Active Motif |

Supplementary Data 1. ROC analysis evaluating the predictive performance of ATAC-seq footprints for *Mef2a* based on the footprint bottom height.

| Footprint bottom height | AUC (Area Under the Curve) | Std. Error | 95% confidence interval | False positive | Ture positive | Threshold | Sensitivity % | 95% confidence interval | Specificity % | 95% confidence interval | Likelihood ration | \|Sensitivity − Specificity\| | Youden index | Likelihood ration (Max) |
|---|---|---|---|---|---|---|---|---|---|---|---|---|---|---|
| 100% | 0.8962 | 0.02843 | 0.8405 ～ 0.9520 | 42 | 74 | < 0.3665 | 81.08 | 70.71% ～ 88.38% | 78.57 | 64.06% ～ 88.29% | 3.784 | 2.51 | 0.5965 | 20.43 |
| 80.0% | 0.9077 | 0.02812 | 0.8526 ～ 0.9628 | 42 | 60 | < 0.3665 | 81.67 | 70.08% ～ 89.44% | 80.95 | 66.70% ～ 90.02% | 4.288 | 0.72 | 0.6262 | 11.55 |
| 60.0% | 0.9167 | 0.02766 | 0.8624 ～ 0.9709 | 42 | 48 | < 0.3665 | 83.33 | 70.42% ～ 91.30% | 80.95 | 66.70% ～ 90.02% | 4.375 | 2.38 | 0.6428 | 21 |
| 50.0% | 0.9252 | 0.02642 | 0.8735 ～ 0.9770 | 42 | 43 | < 0.4500 | 88.37 | 75.52% ～ 94.93% | 78.57 | 64.06% ～ 88.29% | 4.124 | 9.8 | 0.6694 | 9.116 |
| 48.9% | 0.9211 | 0.02773 | 0.8668 ～ 0.9755 | 42 | 40 | < 0.4500 | 87.5 | 73.89% ～ 94.54% | 78.57 | 64.06% ～ 88.29% | 4.083 | 8.93 | 0.6607 | 12.08 |
| 47.5% | 0.92 | 0.02807 | 0.8650 ～ 0.9750 | 42 | 39 | < 0.4500 | 87.18 | 73.29% ～ 94.40% | 78.57 | 64.06% ～ 88.29% | 4.068 | 8.61 | 0.6575 | 21.54 |
| 46.2% | 0.9141 | 0.0299 | 0.8555 ～ 0.9728 | 40 | 38 | < 0.4500 | 86.84 | 72.67% ～ 94.25% | 77.5 | 62.50% ～ 87.68% | 3.86 | 9.34 | 0.6434 | 5.684 |
| 45.0% | 0.9209 | 0.02861 | 0.8649 ～ 0.9770 | 40 | 37 | < 0.4500 | 89.19 | 75.29% ～ 95.71% | 77.5 | 62.50% ～ 87.68% | 3.964 | 11.69 | 0.6669 | 5.838 |
| 42.4% | 0.9285 | 0.02808 | 0.8734 ～ 0.9835 | 39 | 31 | < 0.4500 | 77.42 | 60.19% ～ 88.60% | 87.18 | 73.29% ～ 94.40% | 6.039 | 9.76 | 0.646 | 8.806 |
| 40.8% | 0.9291 | 0.02811 | 0.8740 ～ 0.9842 | 39 | 30 | < 0.6000 | 90 | 74.38% ～ 96.54% | 76.92 | 61.66% ～ 87.35% | 3.9 | 13.08 | 0.6692 | 12.35 |
| 30.0% | 0.9377 | 0.02898 | 0.8809 ～ 0.9945 | 35 | 22 | < 0.6390 | 90.91 | 72.19% ～ 98.38% | 80 | 64.11% ～ 89.96% | 4.545 | 10.91 | 0.7091 | 20.68 |
| 20.0% | 0.9473 | 0.02923 | 0.8900 ～ 1.000 | 32 | 16 | < 0.7085 | 87.5 | 63.98% ～ 97.78% | 84.38 | 68.25% ～ 93.14% | 5.6 | 3.12 | 0.7188 | 20 |

Supplementary Data 2. ROC analysis evaluating the predictive performance of ATAC-seq footprints for *Nkx2.5* based on the footprint bottom height.

| Footprint bottom height | AUC (Area Under the Curve) | Std. Error | 95% confidence interval | False positive | Ture positive | Threshold | Sensitivity % | 95% confidence interval | Specificity % | 95% confidence interval | Likelihood ration | \|Sensitivity - Specificity\| | Youden index | Likelihood ration (Max) |
|---|---|---|---|---|---|---|---|---|---|---|---|---|---|---|
| 100% | 0.6814 | 0.02968 | 0.6232 ～ 0.7396 | 111 | 336 | < 0.2565 | 63.99 | 58.72% ～ 68.94% | 63.06 | 53.79% ～ 71.46% | 1.732 | 0.93 | 0.2705 | 2.343 |
| 80.0% | 0.6963 | 0.03006 | 0.6374 ～ 0.7552 | 110 | 268 | < 0.2970 | 65.3 | 59.42% ～ 70.75% | 64.55 | 55.25% ～ 72.86% | 1.842 | 0.75 | 0.2985 | 2.612 |
| 60.0% | 0.715 | 0.03093 | 0.6544 ～ 0.7756 | 110 | 199 | < 0.3205 | 64.32 | 57.45% ～ 70.65% | 67.26 | 58.16% ～ 75.22% | 1.964 | 2.94 | 0.3158 | 2.633 |
| 50.0% | 0.7328 | 0.03293 | 0.6683 ～ 0.7974 | 102 | 154 | < 0.3485 | 73.38 | 65.89% ～ 79.73% | 61.76 | 52.07% ～ 70.61% | 1.919 | 11.62 | 0.3514 | 2.529 |
| 48.9% | 0.7388 | 0.0328 | 0.6745 ～ 0.8030 | 102 | 150 | < 0.3430 | 66.67 | 58.79% ～ 73.71% | 67.65 | 58.07% ～ 75.94% | 2.061 | 0.98 | 0.3432 | 2.596 |
| 47.5% | 0.7399 | 0.03301 | 0.6752 ～ 0.8046 | 101 | 146 | < 0.3430 | 66.44 | 58.44% ～ 73.59% | 68.32 | 58.71% ～ 76.58% | 2.097 | 1.88 | 0.3476 | 2.767 |
| 46.2% | 0.7339 | 0.03359 | 0.6680 ～ 0.7997 | 99 | 141 | < 0.3430 | 65.96 | 57.81% ～ 73.26% | 67.68 | 57.95% ～ 76.08% | 2.041 | 1.72 | 0.3364 | 2.809 |
| 45.0% | 0.7255 | 0.03418 | 0.6585 ～ 0.7924 | 97 | 138 | < 0.3550 | 66.67 | 58.44% ～ 73.99% | 64.95 | 55.05% ～ 73.71% | 1.902 | 1.72 | 0.3162 | 2.812 |
| 42.4% | 0.7267 | 0.03456 | 0.6590 ～ 0.7944 | 96 | 131 | < 0.3550 | 67.94 | 59.53% ～ 75.32% | 64.58 | 54.62% ～ 73.42% | 1.918 | 3.36 | 0.3252 | 2.931 |
| 40.8% | 0.7327 | 0.03465 | 0.6648 ～ 0.8006 | 95 | 127 | < 0.3660 | 68.5 | 59.98% ～ 75.94% | 65.26 | 55.26% ～ 74.08% | 1.972 | 3.24 | 0.3376 | 2.992 |
| 30.0% | 0.7586 | 0.03665 | 0.6868 ～ 0.8304 | 86 | 86 | < 0.4085 | 68.6 | 58.18% ～ 77.44% | 68.6 | 58.18% ～ 77.44% | 2.185 | 0 | 0.372 | 3 |
| 20.0% | 0.8332 | 0.03536 | 0.7639 ～ 0.9025 | 73 | 57 | < 0.3485 | 75.44 | 62.90% ～ 84.77% | 75.34 | 64.36% ～ 83.80% | 3.059 | 0.1 | 0.5078 | 3.842 |

Supplementary Data 3. ROC analysis evaluating the predictive performance of ATAC-seq footprints for *Tead1* based on the footprint bottom height.

| Footprint bottom height | AUC (Area Under the Curve) | Std. Error | 95% confidence interval | False positive | Ture positive | Threshold | Sensitivity % | 95% confidence interval | Specificity % | 95% confidence interval | Likelihood ration | \|Sensitivity − Specificity\| | Youden index | Likelihood ration (Max) |
|---|---|---|---|---|---|---|---|---|---|---|---|---|---|---|
| 100% | 0.847 | 0.02995 | 0.7883 ～ 0.9057 | 56 | 142 | < 0.2320 | 75.35 | 67.66% ～ 81.71% | 75 | 62.31% ～ 84.48% | 3.014 | 0.35 | 0.5035 | 19.72 |
| 80.0% | 0.8573 | 0.03114 | 0.7963 ～ 0.9183 | 56 | 107 | < 0.2450 | 76.64 | 67.78% ～ 83.64% | 75 | 62.31% ～ 84.48% | 3.065 | 1.64 | 0.5164 | 17.79 |
| 60.0% | 0.8551 | 0.033 | 0.7904 ～ 0.9197 | 54 | 81 | < 0.3095 | 77.78 | 67.58% ～ 85.46% | 74.07 | 61.07% ～ 83.88% | 3 | 3.71 | 0.5185 | 16.67 |
| 50.0% | 0.8665 | 0.03281 | 0.8022 ～ 0.9308 | 52 | 71 | < 0.3095 | 83.1 | 72.74% ～ 90.06% | 73.08 | 59.75% ～ 83.23% | 3.087 | 10.02 | 0.5618 | 16.85 |
| 48.9% | 0.8718 | 0.03218 | 0.8087 ～ 0.9349 | 52 | 69 | < 0.3165 | 84.06 | 73.67% ～ 90.86% | 73.08 | 59.75% ～ 83.23% | 3.122 | 10.98 | 0.5714 | 17.33 |
| 47.5% | 0.8751 | 0.03205 | 0.8123 ～ 0.9380 | 52 | 67 | < 0.3165 | 85.07 | 74.66% ～ 91.69% | 73.08 | 59.75% ～ 83.23% | 3.16 | 11.99 | 0.5815 | 17.07 |
| 46.2% | 0.876 | 0.03292 | 0.8115 ～ 0.9405 | 50 | 63 | < 0.3165 | 87.3 | 76.89% ～ 93.42% | 72 | 58.33% ～ 82.53% | 3.118 | 15.3 | 0.593 | 15.87 |
| 45.0% | 0.874 | 0.03338 | 0.8086 ～ 0.9395 | 50 | 62 | < 0.3165 | 87.1 | 76.55% ～ 93.31% | 72 | 58.33% ～ 82.53% | 3.111 | 15.1 | 0.591 | 15.32 |
| 42.4% | 0.8705 | 0.03471 | 0.8025 ～ 0.9386 | 50 | 55 | < 0.3165 | 87.27 | 75.98% ～ 93.70% | 72 | 58.33% ～ 82.53% | 3.117 | 15.27 | 0.5927 | 14.55 |
| 40.8% | 0.8732 | 0.03537 | 0.8039 ～ 0.9425 | 48 | 52 | < 0.3165 | 90.38 | 79.39% ～ 95.82% | 70.83 | 56.82% ～ 81.76% | 3.099 | 19.55 | 0.6121 | 13.85 |
| 30.0% | 0.8661 | 0.03933 | 0.7890 ～ 0.9432 | 45 | 40 | < 0.3020 | 57.5 | 42.20% ～ 71.49% | 84.44 | 71.22% ～ 92.25% | 3.696 | 26.94 | 0.4194 | 10.13 |
| 20.0% | 0.8758 | 0.04092 | 0.7955 ～ 0.9560 | 40 | 33 | < 0.2970 | 60.61 | 43.68% ～ 75.32% | 85 | 70.93% ～ 92.94% | 4.04 | 24.39 | 0.4561 | 8.485 |

Supplementary Data4. Genomic Datasets

| Datasets | Data Type | Cell or Tissue type | GEO ID | Citation |
|---|---|---|---|---|
| WT-B6.trim.proper_pairs.rmdup.bam | ATAC-seq | Mice Heart left-ventricle tissue [mm10] | | This study |
| SRR8335343.fastq.gz | ChIP-seq | Mice Heart left-ventricle tissue [mm9] | GSE124008 | PMID: 31659164 |
| SRR8335344.fastq.gz | ChIP-seq | Mice Heart left-ventricle tissue [mm9] | GSE124008 | PMID: 31659164 |
| SRR8335345.fastq.gz | ChIP-seq | Mice Heart left-ventricle tissue [mm9] | GSE124008 | PMID: 31659164 |
| SRR8335346.fastq.gz | ChIP-seq | Mice Heart left-ventricle tissue [mm9] | GSE124008 | PMID: 31659164 |
| SRR8335347.fastq.gz | ChIP-seq | Mice Heart left-ventricle tissue [mm9] | GSE124008 | PMID: 31659164 |
| SRR8335348.fastq.gz | ChIP-seq | Mice Heart left-ventricle tissue [mm9] | GSE124008 | PMID: 31659164 |
| SRR8335355.fastq.gz | ChIP-seq | Mice Heart left-ventricle tissue [mm9] | GSE124008 | PMID: 31659164 |
| SRR8335356.fastq.gz | ChIP-seq | Mice Heart left-ventricle tissue [mm9] | GSE124008 | PMID: 31659164 |
| SRR8335357.fastq.gz | ChIP-seq | Mice Heart left-ventricle tissue [mm9] | GSE124008 | PMID: 31659164 |
| SRR8335358.fastq.gz | ChIP-seq | Mice Heart left-ventricle tissue [mm9] | GSE124008 | PMID: 31659164 |
| SRR489735.fastq.gz | ChIP-seq | Mice Heart tissue [mm9] | GSE49847 | PMID: 25409824 |
| SRR489736.fastq.gz | ChIP-seq | Mice Heart tissue [mm9] | GSE49847 | PMID: 25409824 |
| SRR15558986.fastq.gz | CUT&RUN | CD34+ cells derived from fetal liver [hg38] | GSE182530 | PMID: 35941187 |
| SRR15558987.fastq.gz | CUT&RUN | CD34+ cells derived from fetal liver [hg38] | GSE182530 | PMID: 35941187 |
| SRR15558988.fastq.gz | CUT&RUN | CD34+ cells derived from adult peripheral blood [hg38] | GSE182530 | PMID: 35941187 |
| SRR15558989.fastq.gz | CUT&RUN | CD34+ cells derived from adult peripheral blood [hg38] | GSE182530 | PMID: 35941187 |
| SRR15558990.fastq.gz | ATAC-seq | CD34+ cells derived from fetal liver [hg38] | GSE173582 | PMID: 35941187 |
| SRR15558991.fastq.gz | ATAC-seq | CD34+ cells derived from fetal liver [hg38] | GSE173582 | PMID: 35941187 |
| SRR15558992.fastq.gz | ATAC-seq | CD34+ cells derived from adult peripheral blood [hg38] | GSE173582 | PMID: 35941187 |
| SRR15558993.fastq.gz | ATAC-seq | CD34+ cells derived from adult peripheral blood [hg38] | GSE173582 | PMID: 35941187 |
| SRR9044485.fastq.gz | ChIP-seq | CD34+ cells Hematopoietic stem/progenitor cells [hg19] | GSE131052 | PMID: 33512425 |
| SRR9044486.fastq.gz | ChIP-seq | CD34+ cells Hematopoietic stem/progenitor cells [hg19] | GSE131052 | PMID: 33512425 |

### Supplementary information

#### Online methods

##### *Study approval*

All animal experiments were performed following the Guide for the Care and Use of Laboratory Animals (NIH Publication, 8th Edition, 2011) and were approved by the Animal Care and Use Committee of the Osaka University Graduate School of Medicine.

##### *Reagents and antibodies*

All reagents and antibodies used in this study are listed in Supplementary Table 1.

##### *Cardiomyocyte nuclei isolation*

Nuclei were isolated from snap-frozen left ventricular (LV) tissue as previously described with minor modifications[1]. Briefly, 100 mg of frozen LV tissue was minced into small pieces in 3 mL of ice-cold lysis buffer (10 mM Tris-HCl [pH 7.6], 5 mM $CaCl_2$, 2 mM EDTA, 0.5 mM EGTA, 1 mM DTT, 3 mM MgAc, and 0.4% Triton X-100). Tissue was homogenized on ice using a 7 mL glass douncer (30 strokes with a tight pestle). The homogenate was filtered through a 40 µm cell strainer (Falcon, Corning, NY, USA) and centrifuged at 500 × *g* for 5 min at 4 °C to remove cell debris. The nuclear pellet was resuspended in 500 µL of staining buffer (5% BSA, 0.2% Igepal CA-630 in PBS) and incubated with rabbit anti–PCM-1 antibody (1:1,000, Sigma-Aldrich, St. Louis, MO, USA; HPA023370) for 1 hour at 4 °C. After washing, the nuclei were incubated with Alexa Fluor 488–conjugated secondary antibody (1:500, Invitrogen, Carlsbad, CA, USA; A-11034) for 30 minutes at 4 °C. Following a second centrifugation step (500 × *g*, 5 min, 4 °C), the nuclear pellet was resuspended in staining buffer containing DAPI (1:1,000, DOJINDO, Kumamoto, Japan) for nuclear labeling. Nuclei were then sorted using a BD FACSAria II cell sorter (BD Biosciences, San Jose, CA, USA).

***Assay for transposase-accessible chromatin with sequencing (ATAC-seq)***

ATAC-seq was performed using the ATAC-seq Kit (Active Motif, Carlsbad, CA, USA) according to the manufacturer's instructions. Briefly, 50,000 sorted cardiomyocyte nuclei were subjected to tagmentation, followed by PCR amplification using indexed i7 and i5 primers provided in the kit. Libraries were then sequenced using standard paired-end reads (2 × 150 bp) on either a HiSeq X or NovaSeq X Plus platform (Illumina, Hayward, CA, USA). After sequencing, raw reads were demultiplexed, adapter-trimmed, quality-filtered, and aligned to the appropriate reference genome (mm10 for mouse).

***ATAC-seq, ChIP-seq, CUT&RUN, and CUT&Tag data processing and normalization***

Raw sequencing data were initially assessed for quality using FastQC (v0.11.9), followed by adapter trimming with fastp (v0.23.2) using default parameters. Trimmed paired-end reads were aligned to the appropriate reference genome—GRCh38_noalt_as for human, mm10 for mouse—using Bowtie2 (v2.4.5) with the --no-mixed, --no-discordant, and -X 2000 options. SAM files were filtered to remove unmapped reads (-F 0x4) and converted to sorted BAM format using SAMtools (v1.15.1). Only properly paired reads (-f 0x2) were retained, and PCR duplicates were removed using Picard MarkDuplicates (v2.27.4) with REMOVE_DUPLICATES=true. Insert size distributions were calculated with Picard CollectInsertSizeMetrics to assess library quality. Peak calling was performed with MACS2 (v2.2.7.1) using the following parameters: --nomodel, --shift -50, --extsize 100, with genome size specified as hs or mm depending on the species. No control datasets were used. To generate normalized coverage tracks, deepTools (v3.5.1) bamCoverage was applied with the --normalizeUsing CPM and --of bigwig options. These BigWig files were used for visualization in IGV (v2.15.1). All processing steps were executed in a conda-based reproducible environment. Data analysis focused on identifying TF-binding sites through high-resolution footprinting and motif enrichment within accessible chromatin regions, as detailed throughout the main text.

##### *Extraction of high-confidence ChIP-seq peak regions*

High-confidence ChIP-seq peak regions were defined to evaluate motif detection performance. ChIP-seq peaks for each TF (MEF2A, NKX2.5, and TEAD1) were called using MACS2 (v2.2.7.1) with two biological replicates per factor. From each replicate, the top 100 peaks were selected based on $-\log_{10}$(q-value), and the overlapping peaks between the two replicates were identified. This resulted in 77 reproducible peaks for MEF2A, 36 for NKX2.5, and 70 for TEAD1, which were used for downstream motif analysis and footprint validation (Fig. 2).

##### *Identification of heart-specific genes*

To evaluate motif detection performance in a tissue-specific context, we focused on heart-specific genes (Fig. 4). A total of 100 heart-specific genes were selected based on RNA-seq data from the GTEx v10 dataset (https://gtexportal.org), specifically the "Heart - Left Ventricle" tissue. Genes with an average transcript per million (TPM) value ≥300 were retained as heart-specific.

##### *Motif enrichment analysis using HOMER*

Motif enrichment analyses were performed using HOMER (v4.11, http://homer.ucsd.edu/homer/) to identify TF-binding motifs enriched in the input genomic regions. The findMotifsGenome.pl script was applied to ChIP-seq peak regions or ATAC-seq-defined footprints with the corresponding genome build (e.g., mm10 for mouse, hg38 for human), using the following parameters unless otherwise stated: -size given to use the exact region length, and -mask to exclude repetitive sequences. HOMER performs both de novo motif discovery and known motif enrichment analyses. Background regions were automatically selected by HOMER to match GC content and region size, unless otherwise specified via the -bg option. For each motif, enrichment p-values were calculated using the binomial test, and motifs with significant enrichment ($P < 0.05$) were reported. Known motif enrichment results were retrieved from HOMER's internal database of curated TF motifs. In analyses requiring precise motif instance localization, the -find <motif file> option was used to report the genomic positions of matching motifs relative to the input regions. Motif length parameters (e.g., -len 8,10,12) were selected depending on the expected binding site characteristics. All motif analysis results were summarized in HTML-formatted reports and text tables provided in HOMER's output directories.

***Footprint-based motif analysis using TOBIAS***

Footprint-based TF-binding analysis was performed using TOBIAS (v0.12.10) (https://github.com/loosolab/TOBIAS), following the standard three-step pipeline: ATACorrect, FootprintScores, and BINDetect. First, Tn5 insertion bias in ATAC-seq BAM files was corrected using ATACorrect, with the corresponding genome build (mm10 for mouse or hg38 for human) and narrowPeak files from MACS2 peak calling. The corrected bigWig files were then used in FootprintScores to calculate nucleotide-resolution footprint scores across filtered peak regions. Finally, TF-binding site prediction was performed using BINDetect, which scans for motif matches using the JASPAR 2024 CORE vertebrate motif database in MEME format, and classifies each motif occurrence as bound or unbound based on the local footprint signal and accessibility profile. All results, including motif enrichment scores, classification status, and genomic coordinates, were stored in plain-text summary tables generated by TOBIAS.

***The HiFAA framework***

The HiFAA framework is split into three parts (Fig. 1B). The first part performs steps 1 and 2. Finally, in step 3, the exact steps of a HiFAA framework are carried out according to the supplied data. A representative framework run is given in the repository.

*Step 1: Identification of TADs encompassing target genes*

To define the chromatin domain context surrounding selected target genes, we identified TADs based on TF-binding boundaries and gene annotations. The process consisted of three distinct stages: (i) selection of target genes, (ii) TAD boundary definition using CTCF ChIP-seq peaks, and (iii) base-pair resolution coverage extraction from ATAC-seq data.

Step 1 (i): selection of target genes

A list of target genes was prepared based on prior expression profiling, with threshold-based filtering applied to select highly-expressed genes (e.g., TPM >300). The corresponding genomic coordinates—chromosome number, transcription start site (TSS), and transcription end site (TES)—were retrieved by matching gene symbols to reference gene annotations (RefSeq) using the UCSC Genome Browser database for the specific genome assembly (e.g., mm10, hg38). This allowed accurate localization of each gene within the genome.

Step 1 (ii): TAD boundary definition using CTCF ChIP-seq peaks

To define TAD boundaries, we utilized publicly available ChIP-seq data for the architectural protein CTCF (e.g., ENCODE dataset). Peak calling was performed using MACS2 (v2.2.7) with the following parameters. Resulting CTCF peaks were defined by their summit positions. For each target gene, the TAD was defined as the genomic interval bounded by the nearest upstream and downstream CTCF peaks, located within a fixed window (±500 kb) from the gene body. If either flank lacked a CTCF peak within the window, the gene was excluded from downstream analysis to ensure unambiguous domain assignment. Formally, for a gene $G$ with genomic coordinates $[x_{\mathrm{TSS}}, x_{\mathrm{TES}}]$, the TAD region was defined as:

$$\mathrm{TAD}G = \left[max\{x_i \in P | x_i < x_{\mathrm{TSS}}\}, min\{x_j \in P \middle| x_j > x_{\mathrm{TES}}\} \right]$$

where $P$ is the set of CTCF peak summits.

Step 1 (iii): Base-pair resolution coverage extraction from ATAC-seq data

To characterize the chromatin accessibility landscape within each TAD, ATAC-seq data were used to extract per-base read coverage. BAM files were filtered to include only properly paired reads and to exclude PCR duplicates. For each TAD region, 1-bp resolution read coverage was calculated using pysam (v0.22.0) and stored in CSV format. The read coverage at position $x$ was defined as:

$$\mathrm{Coverage}(x) = \sum_{r \in R} 1_{x \in r}$$

where $R$ is the set of filtered paired-end ATAC-seq reads, and $1_{x \in r}$ is an indicator function that equals 1 if position $x$ is covered by read $r$, and 0 otherwise. This base-pair resolution coverage information served as the input for subsequent footprint detection and motif ranking.

*Step 2: Footprint shape definition and pseudo-read transformation*

Chromatin accessibility footprints, which represent short protected regions of DNA bound by TFs, are characterized by localized depletions in ATAC-seq signal flanked by enriched regions of transposase cutting activity. To computationally define such structures, we developed a multi-stage process that extracts, refines, and encodes the signal shape of candidate footprints from aligned ATAC-seq data with single base-pair resolution. The process consisted of six distinct stages: (i) per-base signal extraction, (ii) discretized region segmentation, (iii) read-count hierarchy transformation, (iv) noise filtering and thresholding, (v) peak-to-region integration, and (vi) pseudo-read matrix generation.

Step 2 (i): per-base signal extraction

Using the BAM files from ATAC-seq data (see step 1), all aligned read midpoints were counted on a base-pair level. For each genomic region $R = [s, e]$, we constructed a vector:

$$\mathrm{v}_R = [c_1, c_2, \cdots, c_L],\ \ L = e - s + 1$$

where $c_i \in \mathbb{Z}_{\geq 0}$ represents the number of transposase cut sites mapped at the $i$-th base within the region $R$. This signal vector $\mathrm{v}_R$ was stored as a CSV-formatted file for each genomic interval, enabling efficient downstream parsing.

Step 2 (ii): discrete region segmentation

The signal vector $\mathrm{v}_R$ was converted into discrete regions by grouping adjacent positions with identical read counts. For a given read-count value $k$, a set of positions was defined as:

$$S_k = \{x \in [1, L] | c_x = k\}$$

This set was then split into maximal continuous subsequences $[a_j, b_j]$ satisfying:

$$\forall_i \in [a_j, b_j], c_i = k \ \text{ and } \ c_{a_j} \neq k, c_{b_{j+1}} \neq k$$

These subsequences represent plateaus of uniform signal intensity within the footprint landscape and form the basic building blocks for further shape definition.

Step 2 (iii): read-count hierarchy transformation

To enrich the detection of high-confidence footprint features, we introduced a cumulative signal aggregation strategy. For a given threshold k, we defined a cumulative binary vector $\tilde{\mathrm{v}}_k$ such that:

$$\tilde{v}_i^{(k)} = \begin{cases} 1, & if\ c_i \geq k \\ 0, & otherwise \end{cases}$$

This transformation allowed us to represent, for each threshold $k$, a signal trace of accessible bases that met or exceeded the threshold. This step implicitly constructed a hierarchical view of accessibility, where higher $k$ values indicate progressively more confident footprints. In practice, thresholds from $k = 1$ up to the maximal observed read count within the region were iteratively applied, generating a series of binary matrices corresponding to increasing stringency.

Step 2 (iv): noise filtering and thresholding

To reduce signal fragmentation due to technical noise or background accessibility, we applied a minimum read-count threshold T to exclude low-confidence positions. This threshold was empirically determined based on the global read-count distribution (typically $T = 3$ unless otherwise noted). All contiguous intervals satisfying $c_i \geq T$ were merged into "active" regions. For each such merged interval $M = [a, b]$, we identified local maxima ("peak tops") satisfying:

$$c_i > c_{i-1} \text{ and } c_i > c_{i+1},\ \forall_i \in [a+1, b-1]$$

Each peak $P_j$ was then annotated with its summit position $S_j$, height $h_j = c_{S_j}$, and local support width, initially defined as the full width at half maximum (FWHM):

$$\text{FWHM}(P_j) = \max\left\{r - l + 1 \middle| c_l, c_r \geq \frac{h_j}{2}, l < S_j < r\right\}$$

Step 2 (v): peak-to-region integration

In cases where multiple peaks occurred within a single region, we performed a comparative prominence analysis to distinguish independent events from artifacts or shoulders of broader peaks. Given two neighboring peaks $P_a$ and $P_b$ with summits $S_a$, $S_b$ and heights $h_a$, $h_b$, we computed the local valley depth $h_v$ between them and defined the prominence ratio as:

$$\text{Prominence}(P_a, P_b) = \frac{\min(h_a, h_b) - h_v}{\min(h_a, h_b)}$$

Only peak pairs with $\text{Prominence} \geq \theta$ (e.g., $\theta = 0.3$) were retained as distinct, otherwise they were merged. Additionally, overlapping footprint candidates were sorted by descending read count, and non-top peaks that overlapped with higher-ranked peaks were clipped or removed depending on their residual prominence.

Step 2 (vi): pseudo-read matrix generation

For each finalized footprint peak $P_j$, a pseudo-read profile was extracted within a symmetric window of size $2\omega + 1$ centered on the summit $S_j$, with $\omega = 100\,\text{bp}$ unless otherwise specified:

$$f_j = \left[c_{s_j-\omega}, \cdots, c_{s_j}, \cdots, c_{s_j+\omega}\right]$$

These vectors $f_j$ were stored in matrix form $F \in \mathbb{Z}_{\geq 0}^{N\times(2\omega+1)}$, where $N$ is the total number of footprint peaks identified in the dataset. Each row represented the accessibility shape around a candidate binding site and served as input to downstream motif matching or clustering algorithms. To preserve positional alignment for motif analysis, strand orientation was maintained. For reverse-strand motifs, the accessibility profile $f_j$ was reversed to align upstream and downstream features consistently.

*Step 3: Quantitative ranking of footprint activity*

To prioritize chromatin accessibility footprints with the highest likelihood of representing true TF-binding events, we implemented a quantitative ranking framework that integrates footprint geometry with absolute accessibility features. This stage converts the pseudo-read matrices generated in step 2 into high-confidence, rank-ordered footprint calls for downstream motif inference and TF activity modeling. The process consisted of four distinct stages: (i) central footprint bottom height extraction, (ii) global percentile-based filtering, (iii) footprint coordinate export, and (iv) motif enrichment analysis with HOMER.

Step 3 (i): Central footprint bottom height extraction

For each finalized footprint peak $P_j$, a pseudo-read profile was extracted within a symmetric window of size $2\omega + 1$ centered on its summit position $S_j$:

$$f_j = \left[c_{j,s_j-\omega}, \cdots, c_{j,s_j}, \cdots, c_{j,s_j+\omega}\right] \in \mathbb{Z}_{\geq 0}^{2\omega+1}$$

where $S_j$ denotes the summit (i.e., the base with the minimal accessibility signal within the footprint), and $\omega$ is the half-window size (default: 100 bp). From this profile, we extracted the central read count $c_{j,s_j}$, which directly quantifies the number of ATAC-seq insertions at the footprint bottom. In the HiFAA system, higher values of $c_{j,s_j}$ are interpreted as more robust footprints, reflecting confident signal presence rather than depletion. Unlike relative depletion metrics that rely on flanking normalization (e.g.,

$$D_j = \frac{\mu_{\text{flank}} - c_{j,s_j}}{\mu_{\text{flank}} + \epsilon}$$

), HiFAA ranking exclusively uses the absolute footprint bottom height $c_{j,s_j}$. This enabled reduced sensitivity to local background variability and enhanced detection of true TF occupancy signals.

Step 3 (ii): global percentile-based filtering

To retain the most protected and structurally stable footprints, all candidate entries were sorted in descending order by their bottom height $c_{j,s_j}$, and a global percentile cutoff was applied. Let

$\mathcal{P} = \{P_1, P_2, \cdots, P_N,\}$ denote the full set of candidate footprints from step 2. The filtered subset was then defined as:

$$\mathcal{F} = \left\{P_j \in \mathcal{P} \middle| \text{Rank}\left(c_{j,s_j}\right) \leq 0.489 \times N\right\}$$

Here, $\text{Rank}\left(c_{j,s_j}\right)$ refers to the descending order rank of the central read count—i.e., higher counts receive better ranks. The threshold of 48.9% was determined empirically from multiple datasets to optimize sensitivity and specificity in motif enrichment performance (see Extended data Fig. 3, Extended data Fig. 4, Extended data Fig. 5, Supplementary Data 1, Supplementary Data 2, Supplementary Data 3). Only footprint profiles in $\mathcal{F}$ were retained for subsequent analysis.

Step 3 (iii): footprint coordinate export

The genomic coordinates corresponding to the selected footprints $\mathcal{F}$ were exported in BED format, facilitating compatibility with motif discovery tools such as HOMER. Each BED entry included: chromosome, genomic start and end coordinates of the window $\left[s_j - \omega, s_j + \omega\right]$, strand orientation, and an optional score field (e.g., normalized $c_{j,s_j}$). This standardized output enabled genome-wide annotation and cross-sample comparison of candidate binding events.

Step 3 (iv): motif enrichment analysis with HOMER

To assess the sequence-level relevance of the selected footprints, motif enrichment analysis was performed using HOMER (v4.11). The top 48.9% of ranked footprints were converted to BED format, with each region defined as a 47 bp window centered on the footprint summit (±23 bp). These regions were analyzed using the findMotifsGenome.pl script with default parameters for short motifs (-len 8,10,12) against the corresponding reference genome. A matched background set of 47 bp regions was generated automatically by HOMER to control for sequence bias. Significantly enriched motifs (adjusted $p < 10^{-4}$) were used to confirm TF-binding specificity within the prioritized footprints.

***Sensitivity and specificity evaluation of transcription factor binding site (TFBS) predictions***

To compare the TFBS prediction performance of our method (HiFAA) with established tools (TOBIAS and HOMER), we quantitatively evaluated sensitivity and specificity using experimentally validated ChIP-seq peaks as ground truth. Because each method differs in footprint definition and motif annotation and classification, we tailored the definitions of sensitivity and specificity to reflect the logic and output structure of each tool. Sensitivity was defined as the proportion of ChIP-seq peaks that overlap with motif-containing regions predicted by each method. Accordingly, both the numerator and denominator in the sensitivity metric are directly dependent on the total number of ChIP-seq peaks. Specificity, in contrast, was defined as the proportion of motif-containing ATAC-seq footprints that are supported by ChIP-seq evidence. Thus, both the numerator and denominator of the specificity metric are based on the number of ATAC-seq-derived footprint regions carrying motifs. This formulation ensures that each metric reflects the appropriate biological axis: sensitivity measures coverage of true binding events (ChIP-seq peaks), whereas specificity assesses the precision of predicted motif sites within accessible chromatin (ATAC-seq footprints).

*HiFAA*

HiFAA first defines high-confidence footprint regions based on ATAC-seq signal features, and subsequently performs motif scanning within those footprint intervals. Thus, its predictions are limited to motif occurrences located within valid footprints.

For HiFAA, we defined:

• Sensitivity as the proportion of ChIP-seq peaks that intersect with motif-containing footprints:

$$\text{Sensitivity}_{\text{HiFAA}} = \frac{Number\ of\ (\text{Motif}\cap\text{Footprtint}\cap\text{ChIP})}{Number\ of\ (\text{ChIP})}$$

• Specificity as the proportion of motif-containing footprints that overlap ChIP-seq peaks:

$$\text{Specificity}_{\text{HiFAA}} = \frac{Number\ of\ (\text{Motif}\cap\text{Footprtint}\cap\text{ChIP})}{Number\ of\ (\text{Motif}\cap\text{Footprtint})}$$

This definition reflects HiFAA's design: only those footprints that are both motif-positive and ChIP-positive are considered true positives, and all motif-containing footprints are evaluated regardless of ChIP-seq status.

*HOMER*

HOMER was used to perform motif enrichment analysis using a fixed set of ATAC-seq peak regions. These peaks were first identified by MACS2 (q <0.01) from aligned ATAC-seq data, representing regions of chromatin accessibility. The HOMER findMotifsGenome.pl function was then used to identify significantly enriched TF-binding motifs within these peaks. HOMER does not infer footprinting, nor does it classify motif instances as bound or unbound.

Given this structure, we defined sensitivity and specificity as follows:

• Sensitivity was calculated as the proportion of ChIP-seq peaks that overlapped with HOMER-predicted motif instances:

$$\text{Sensitivity}_{\text{HOMER}} = \frac{Number\ of\ (\text{Motif}\cap\text{ChIP})}{Number\ of\ (\text{ChIP})}$$

• Specificity was defined as the proportion of HOMER-predicted motifs that fell within ChIP-seq peaks:

$$\text{Specificity}_{\text{HOMER}} = \frac{Number\ of\ (\text{Motif}\cap\text{ChIP})}{Number\ of\ (\text{Motif})}$$

These metrics enable a motif-level evaluation despite HOMER's lack of accessibility modeling or classification of TF occupancy.

*TOBIAS*

TOBIAS differs fundamentally from HiFAA in that it does not filter footprints prior to motif scanning. Instead, it scans all accessible chromatin regions for motifs and computes footprint scores, subsequently classifying each motif occurrence as bound or unbound based on signal depth and local accessibility. This relative approach enables broader motif evaluation but introduces additional complexity in interpreting predictive performance.

To account for this, we defined:

• Sensitivity as the fraction of ChIP-seq peaks overlapping motif-containing bound footprints:

$$\text{Sensitivity}_{\text{TOBIAS}} = \frac{Number\ of\ (\text{Motif}\cap\text{Bound}\cap\text{ChIP})}{Number\ of\ (\text{ChIP})}$$

• Specificity was defined as the proportion of HOMER-predicted motifs that fell within ChIP-seq peaks:

$$\text{Specificity}_{\text{TOBIAS}} = \frac{Number\ of\ (\text{Motif}\cap\text{Bound}\cap\text{ChIP})}{Number\ of\ (\text{Motif}\cap\text{Bound})}$$

These definitions align with TOBIAS's bound/unbound classification logic and enable consistent comparison with other tools under standardized conditions.

***Balanced performance assessment using Geometric-mean***

To comprehensively evaluate the classification performance of TF-binding site prediction tools, we calculated the Geometric-mean of sensitivity and specificity:

$$\text{Geometric} - \text{mean} = \sqrt{\text{Sensitivity}\times\text{Specificity}}$$

The Geometric-mean metric is widely adopted in imbalanced classification tasks, as it penalizes models that perform well on one metric (e.g., sensitivity) at the expense of the other (e.g., specificity). In the context of transcription factor footprinting, where both the ability to detect true TF-binding sites (sensitivity) and the precision in avoiding false positives (specificity) are critical, Geometric-mean provides a robust, single-value summary that balances these two axes of performance. Unlike F1-score, which requires binary labels (e.g., positive/negative predictions), Geometric-mean can be applied directly to motif-level overlaps with ChIP-seq peaks and does not rely on threshold-based binarization. Thus, it is particularly suited for evaluating footprinting frameworks that produce ranked, signal-based outputs.

***Statistical analysis***

All data are presented as mean ± standard deviation (s.d.) unless otherwise noted. For comparisons between two groups, unpaired two-tailed Student's t-test was used to assess statistical significance. A P value $<0.05$ was considered statistically significant. Statistical tests and data visualization were performed using GraphPad Prism software (MDF; Tokyo, Japan). ROC curve analysis was used to assess the classification performance of ATAC-seq-defined footprints. Footprints overlapping both ChIP-seq peaks and matching TF motifs were considered positive, while those containing motifs but lacking ChIP-seq support were considered negative. ROC curves were generated by varying the threshold of the height of footprint bottom signal, and the AUC was calculated to quantify classification accuracy at different ranking thresholds.

**Data availability**

ChIP-seq datasets for MEF2A, NKX2.5, and TEAD1 in adult mouse heart (P42 ventricle apex) were obtained from GEO under accession GSE124008 (originally aligned to mm9; reprocessed to mm10 for this study)[2]. ChIP-seq data for CTCF in adult mouse heart (8-week-old) was retrieved from GEO under accession GSE49847 (aligned to mm10)[3]. ATAC-seq data from adult mouse heart muscle (8-week-old) were newly generated in this study and are available upon request. These data will be deposited in GEO under the provisional accession GSEXXXXXX (aligned to mm10). GATA1 CUT&RUN and ATAC-seq data for fetal and adult erythroblasts (CD34+ cells derived from fetal liver and adult peripheral blood, both differentiated into the erythroid lineage on day 11) were obtained from GEO under accession numbers GSE182530 and GSE173582 (aligned to hg38)[4]. ChIP-seq data for CTCF in hematopoietic stem/progenitor cells and erythroblasts (12-day erythropoiesis) were obtained from GEO under accession GSE131052 (originally aligned to hg19; reprocessed to hg38)[5]. All external datasets were downloaded as raw FASTQ files and reprocessed using either mm10 (mouse) or hg38 (human) genome assemblies for consistency across analyses. The sequencing data used in this study is shown in Supplementary Data 4. All other data supporting the findings of this study are available from the corresponding author upon reasonable request.

**Code availability**

The HiFAA software is publicly available at GitHub.
